# Informed dimension reduction of clinically-related genome-wide association summary data characterises cross-trait axes of genetic risk

**DOI:** 10.1101/2020.01.14.905869

**Authors:** Oliver S Burren, Guillermo Reales, Limy Wong, John Bowes, James C Lee, Anne Barton, Paul A Lyons, Kenneth GC Smith, Wendy Thomson, Paul DW Kirk, Chris Wallace

## Abstract

Integration of genome-wide association study (GWAS) data has been used to generate new hypotheses of biological mechanism, aetiological relationships between traits, or test causality of one factor for another. However, such approaches have typically been limited to pairwise comparisons of traits. We propose a generally applicable method, that exploits ideas from Bayesian genetic fine mapping to define a “lens” that focuses relevant variants before dimension reduction of a set of related GWAS summary statistics. We applied this technique to immune-mediated diseases, deriving 13 components which summarise the multidimensional patterns of genetic risk. Projection of independent datasets demonstrated the specificity and accuracy of our reduced dimension basis, enabled us to functionally characterise individual components, identify disease-discriminating components and suggest novel associations in rare diseases where classical GWAS approaches are challenging. Our approach summarises the genetic architectures underlying any range of aetiologically-related traits in fewer dimensions, facilitating more nuanced multidimensional comparative analyses.

---

The collected summary data of genome-wide association studies (GWAS) represent, in a compressed form, assays of thousands of phenotypes across millions of common genetic variants. Analysed individually, GWAS have elucidated the polygenic component of common human diseases^1^ and comparative studies of summary GWAS results have highlighted a shared genetic aetiology across different diseases.^2^ However, comprehensive overviews of sharing between multiple diseases are made difficult by the dimension of these statistics (100,000s of SNPs), the complex patterns that exist, and the limitation that while all dimensions carry information about technical differences between studies (DNA storage, processing, and population sampling), only a minority carry detectable information about disease risk. Therefore, such analyses have typically been approached from one of two angles: a variant-by-variant analysis across multiple diseases focusing on individual variants in turn,^3,4^ or pairwise analysis of diseases across multiple variants at a regional or genome-wide level.^5,6^ Both approaches have limitations. Different patterns of sharing identified at different variants make generalisations about inter-disease relationships difficult, while disease-pairwise approaches make comparison of more than two diseases challenging. Thus, a need exists to represent a multi-dimensional view of shared genetic architectures between multiple diseases.

The GWAS approach explicitly accounts for the number of tests (SNPs) by requiring successively larger samples (tens of thousands). Large samples present an insurmountable barrier for rare diseases, where efforts have instead focused on searching for rare variants of high penetrance through whole exome^7^ or whole genome^8,9^ sequencing. Despite this, moderate-sized GWAS-style studies of rare diseases have found both polygenic association with common variants^9,10^ and evidence for differential genetic associations between clinical subtypes of these rare diseases.^11^ Thus, a need also exists to democratise GWAS to less common diseases, which may be possible by considering them in the context of commoner, yet related diseases.

We propose studying multifactorial genetic risks of related diseases in an informed dimension-reduction approach based on matrix decomposition. Matrix decomposition, for example via principal component analysis (PCA), expresses a matrix as the product of two smaller matrices, and has been used extensively in genetics to summarise population structure and address its confounding effects in association studies.^12^ It has also been used to explore structure in genetic association with multiple traits, either from different studies through aggregated signals across SNPs according to physical proximity to genes^13^ or using a linkage disequilibrium (LD) independent subset of SNPs from a single cohort^14^ In either case, the reduced dimensional space was used to explore the same datasets as used to define it, with two implications. First, GWAS summary statistics are a composite of biological signal, technical noise, and sampling variation. Decomposition aims to find axes that maximise variance explained in the input datasets, and cannot distinguish between these three sources of variability. We therefore expect it to magnify technical and random differences as well as biological, a problem related to over-fitting in high-dimensional datasets. Second, there is no treatment of uncertainty in the reduced dimension space, meaning we can measure the distance between diseases, but not test whether that difference is non-zero.

Here we propose augmenting PCA of GWAS summary statistics by a Bayesian shrinkage approach that mitigates overfitting. Our central aim is to define a reduced dimension space, with axes that describe different patterns of genetic susceptibility corresponding to underlying biological risk factors. In a transfer learning paradigm, we can project independent datasets into this space, allowing us to study the distinct and shared genetic contributions to related diseases, and use standard statistical techniques to test for genetic association of rare diseases or genetic differences between disease subtypes. We use immune-mediated diseases (IMD) as a paradigm for a set of traits with established aetiological overlap^2^ to highlight the potential uses of this method.

## Results

### A genetic basis for immune mediated diseases

We used well-powered, publicly available case/control GWAS summary statistics (estimated log odds ratios, 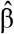) across 13 IMD (Supplementary Table 1) to derive a principal component *basis*. Studies were chosen to balance the competing aims of maximising the number of studies, the number of SNPs common to all studies, and the number of samples in each study (to minimise noise in 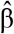). We excluded all variants within the MHC (GRCh37 Chr6:20-40Mb) due to its long and complex LD structure, and because SNPs in the MHC have a profound involvement in IMD susceptibility, and thus the potential to dominate the basis, overwhelming more modest signals. As in conventional PCA, this basis consists of orthogonal principal components (PCs), constructed as linear combinations of input 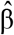, which together provide a lower dimensional representation of genetic associations with IMD.

To ensure the disease-relevance of the basis, we wanted to preferentially use information from truly associated SNPs, while avoiding double counting evidence from SNPs in LD. DeGAs^14^ deals with this by thinning SNPs by LD and hard thresholding, replacing 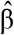 by Z scores, setting these to 0 when the associated p > 0.001. As Z scores are standardised 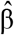, this has the effect of shrinking 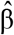 towards 0 when uncertainty is high, such as when allele or disease frequencies are low, which means information from more common diseases will dominate. We chose instead to only partially standardise (for the effects of allele frequency), and to deal with LD and remaining noise simultaneously via shrinkage, adopting ideas from Bayesian fine mapping which jointly models association across neighbouring SNPs. This allowed us to define a continuous weight which adaptively shrinks 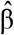 towards 0 (Fig. 1, Methods). Finally, we report projected results as 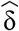, the difference between the projected 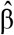 and a projected synthetic control with all entries 0, which allows us to make statistical inference about whether its estimand, δ, differs from control.

**Figure 1:**
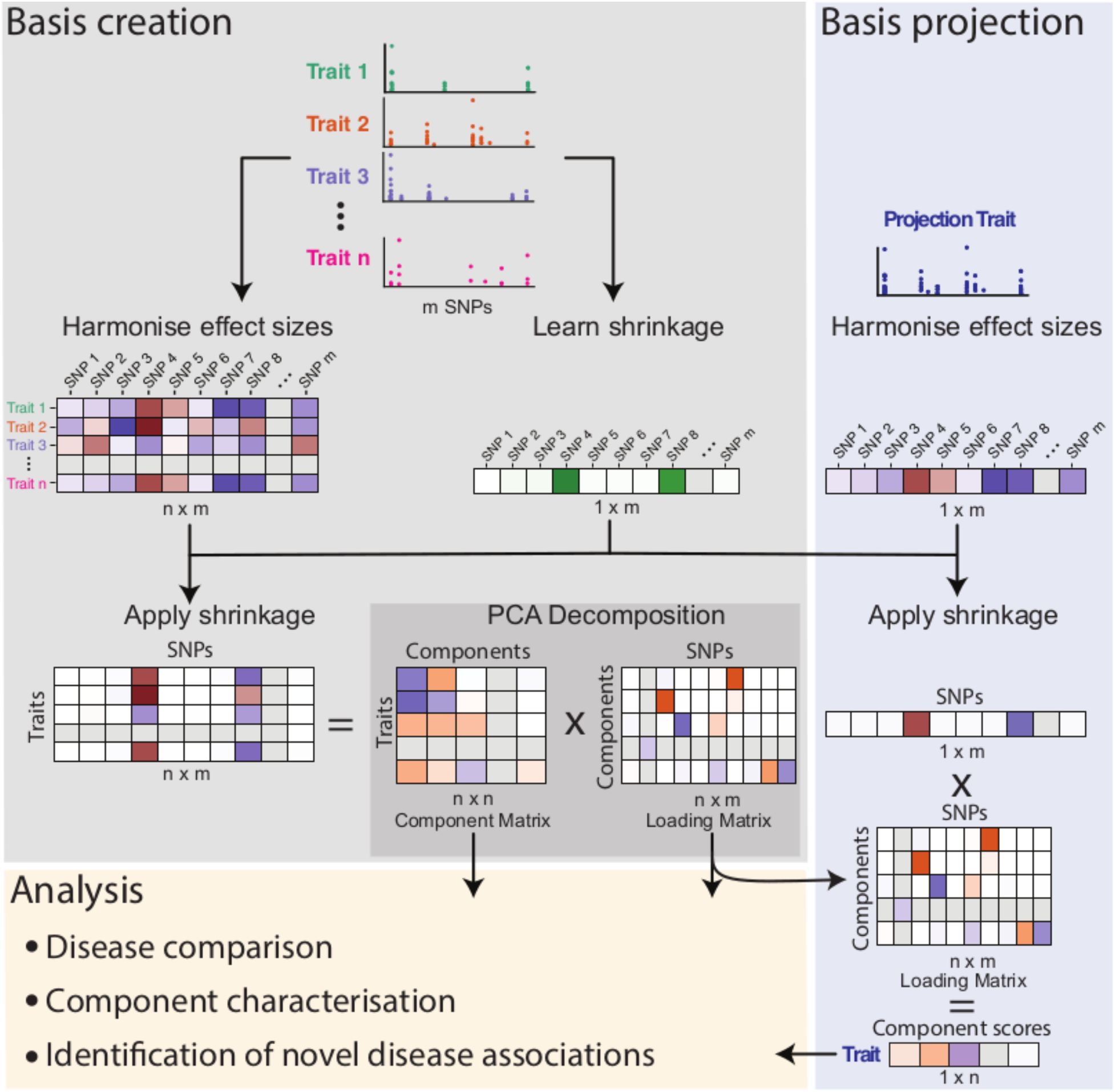
Schematic of basis creation and projection. Basis creation: GWAS summary statistics for related traits are combined to create a matrix, M (n x m), of harmonised effect sizes 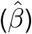 and a learned vector of shrinkage values for each SNP. After multiplying each row of M by the shrinkage vector, PCA is used to decompose M into component and loading matrices. Basis projection: External trait effects are harmonised with respect to the basis, shrinkage applied and the resultant vector is multiplied by the basis loading matrix to obtain component scores which can then be used for further analyses.

To illustrate the importance of our informed shrinkage procedure, we built four bases, with GWAS summary statistics for the 13 IMDs shrunk differently in each case. We assessed their relative performance by projection of matching self-reported diseases (SRD) from UK BioBank (UKBB)^15^ using summary statistics from a compendium provided by the Neale lab [http://www.nealelab.is/uk-biobank/]. While all bases found structure in the input data, in the basis without shrinkage (Fig. 2a), the UKBB SRD clustered with each other rather than their GWAS comparator, suggesting that the structure identified related to between study differences other than disease. In hard-thresholded, LD-thinned bases using either Z scores or 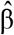 (Supplementary Fig. 1), some of the structure identified was disease-related for the larger GWAS of more common traits (asthma, multiple sclerosis [MS], Crohn’s disease, ulcerative colitis [UC]). In contrast, in the basis created with continuous shrinkage (Fig. 2d), the UKBB SRD clearly clustered with their GWAS comparators, suggesting that the structure captured is disease-relevant, such that UKBB data from relatively infrequent diseases such as type 1 diabetes (T1D) (318 cases) and vitiligo (105 cases) are projected onto the same vectors as their larger comparator GWAS studies. The basis generated is naturally sparse (Supplementary Fig. 2), enabling us to identify 107-373 “driver SNPs” that are required to capture genetic associations on any individual component.

**Figure 2:**
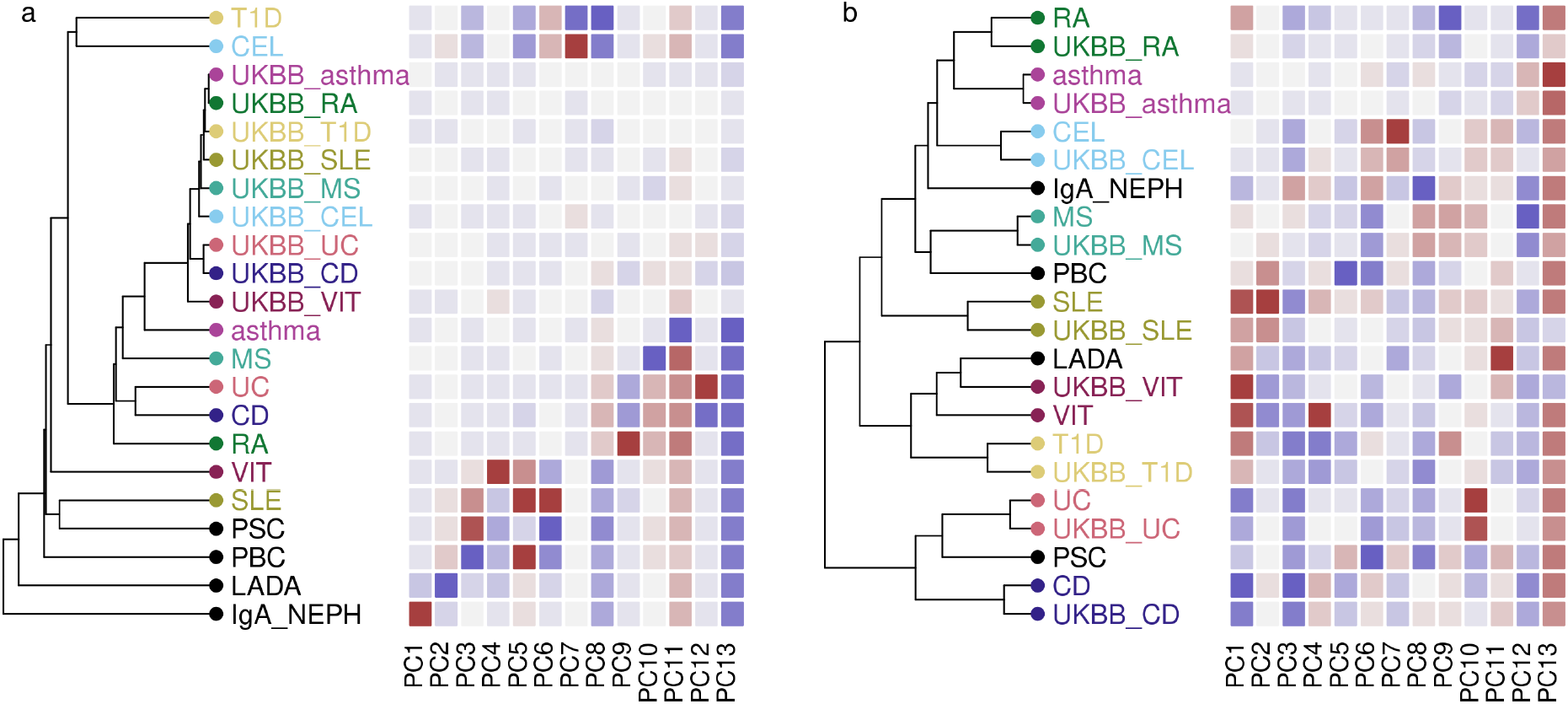
Hierarchical clustering of basis diseases and their UKBB counterparts in **a** unweighted basis constructed using 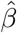 **b** basis constructed using continuous shrinkage applied to 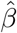. Heatmaps indicate projected 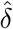 for each disease on each component PC1-PC13, with grey indicating 0 (no difference from control), and darker shades of blue or magenta showing departure from controls in one direction or the other. GWAS datasets: T1D = type 1 diabetes, CEL= celiac disease, asthma, MS =multiple sclerosis, UC =ulcerative colitis, CD = Crohn’s disease, RA =rheumatoid arthritis, VIT =vitiligo, SLE =systemic lupus erythematosus, PSC=primary sclerosing cholangitis, PBC=primary biliary cholangitis, LADA=latent autoimmune diabetes in adults, IgA_NEPH= IgA nephropathy. UKBB_ prefixed diseases correspond to self reported disease status in UK Biobank.

We projected data from three classes of study onto the basis with shrinkage. First, we used all self-reported disease and cancer traits from UKBB to characterise the basis components, to examine specificity to IMD, and to assess power as a function of sample size: case numbers for UKBB self reported IMD range from 41,000 (asthma) to 105 (vitiligo). Second, we used IMD GWAS with smaller sample sizes than used in basis construction, including diseases studied in multiple ancestral backgrounds to explore robustness to ancestry. Third, we used studies of IMD that are too rare or clinically heterogeneous to build large GWAS cohorts.

We also created online tool to allow projection and exploration of additional data into the basis (https://grealesm.shinyapps.io/IMDbasisApp/).

### Genetic analysis in reduced dimensions

Across all 312 projected UKBB traits (Supplementary Table 2), 27 had significantly non-zero 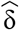 (FDR < 1%). These were overwhelmingly immune-related traits (Supplementary Fig. 3): no significance was observed for traits such as coronary artery disease, stroke, or obstructive sleep apnea, confirming the immune-mediated specificity of our basis. Significant results were detected with as few as 105 cases for vitiligo, emphasising the potential of this approach to unlock the genetics of rare IMD GWAS.

Of 28 traits from target (non-UKBB) IMD GWAS, including JIA, neuromyelitis optica (NMO), vasculitis and their clinical subtypes, 16 were significant (FDR < 1%, Supplementary Table 3, Supplementary Fig. 4-16). We clustered all 28, together with significant UKBB diseases to generate a visual overview of IMD and associated traits (Fig. 3). This highlighted two small groups, inflammatory bowel disease (IBD) and EGPA, and two larger groups, one comprising autoimmune diseases and the other a heterogeneous cluster containing subgroups centred on MS, ankylosing spondylitis (AS), atopy, and traits with only weak or non-significant signals. Notably, three studies of AS all clustered together, despite only one having sufficient sample size for significant results and the three studies representing different ancestries (UK-European, International and Turkish/Iranian). A broader examination of projecting non-European samples found that projections are generally attenuated but consistent with projections of European samples (Supplementary Fig. 17, Supplementary Note).

**Figure 3:**
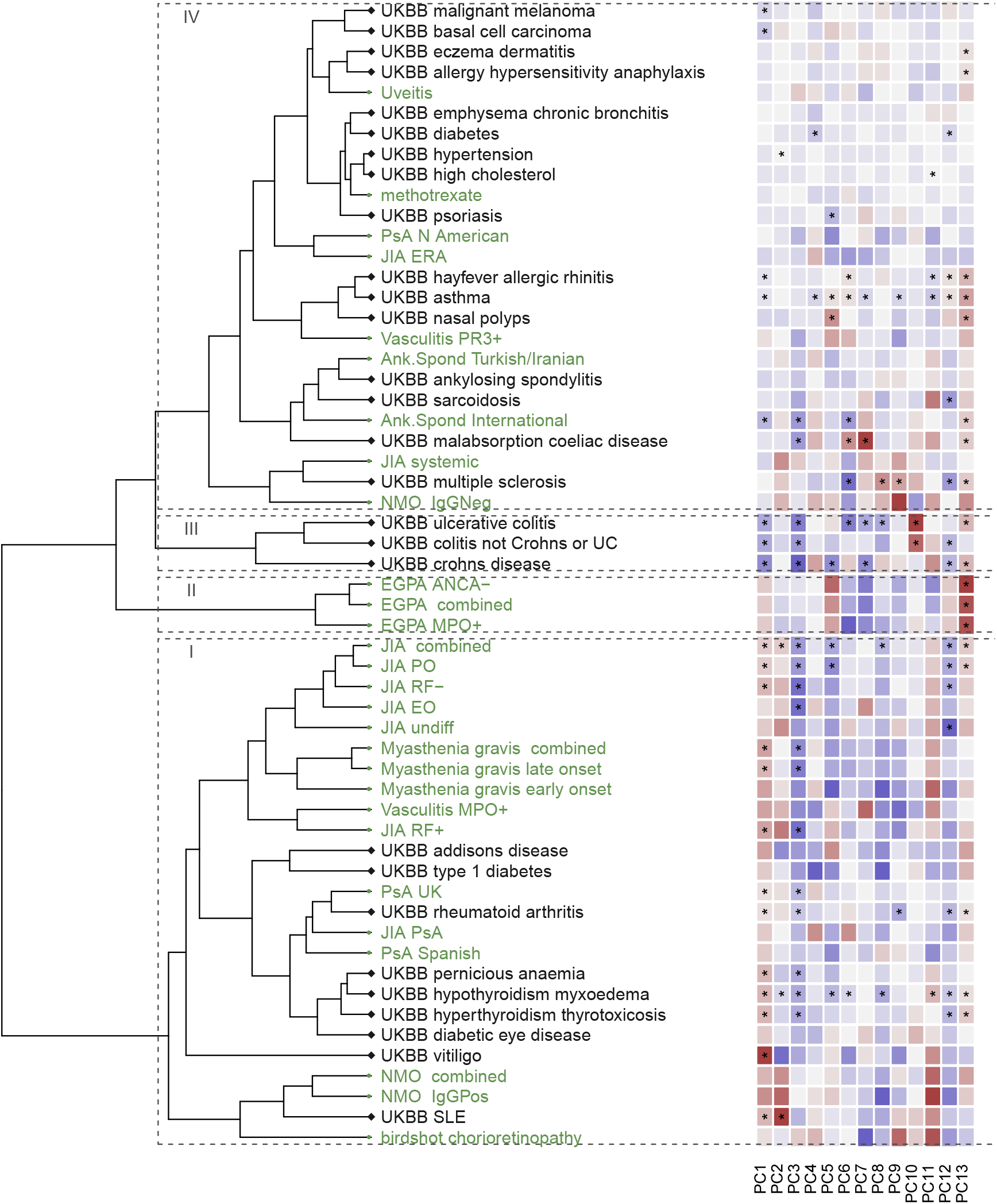
Hierarchical clustering of projected diseases significantly different from control (FDR < 1%) or of small sample size. Coloured labels are used to distinguish UKBB (grey) and other GWAS (green) datasets. Heatmaps indicate delta values for each disease on each component PC1-PC13, with grey indicating 0 (no difference from control), and darker shades of blue or magenta showing departure from controls in one direction or the other. An overlaid * indicates delta was significantly non zero (FDR<0.01). Roman numerals indicate clusters described in the text. Abbreviations: ANCA- = antineutrophil cytoplasmic antibody negative, Ank. Spond = ankylosing spondylitis, EGPA = eosinophilic granulomatosis with polyangiitis, EO = extended oligo, ERA = juvenile enthesitis-related arthritis, IgG- Pos = IgG positive, JIA = juvenile idiopathic arthritis, MPO+ = myeloperoxidase positive NMO = neuromyelitis optica, PO = persistent oligo, PR3+ = proteinase 3 positive, PsA = psoriatic arthritis, RF +/- = polyoligo rheumatoid factor positive/negative, SLE = systemic lupus erythematosus, UC = ulcerative colitis.

We expect that significant results represent a composite of many small effects working in consistent directions. However, false positives could also result if a single SNP with a large weight in the basis is in LD with a SNP with a large effect on the projected trait due to chance. To guard against this, we used Spearman rank correlation which is robust to such outlier observations to test the “consistency” of each projection (Supplementary Note). We found, reassuringly, that increasing deviation from control correlated with increasing consistency amongst disease traits (Supplementary Fig. 18).

Some individual components could be biologically interpreted due to their pattern of disease or other trait associations. **PC1**, which explained the greatest variation in the training datasets, appears to represent an autoimmune/(auto)inflammatory axis^16^, also characterised by whether diseases are considered antibody ‘seropositive’ / ‘seronegative’ (Fig. 4). The exception is vitiligo, in which, despite strong evidence of T cell autoimmunity, autoantibodies are reported but are not consistent features of disease.^17^ Weaker but significant association of Psoriatic arthritis (PsA) among the other seropositive IMD is also consistent with a recent report of novel pathogenic antibodies in PsA.^18^ On the inflammatory/seronegative side, we also saw weaker but still significant signals for atopy, basal cell carcinoma and malignant melanoma. Both malignant melanoma and non-melanoma skin cancer incidence is increased in IBD, but the relative role of treatment or IBD itself in driving this is hard to determine.^19,20^ On the seropositive side, we saw significant results for pernicious anemia, a disease strongly associated with anti-gastric parietal cell and anti-Intrinsic Factor antibodies, as well as with autoimmune thyroiditis, T1D and vitiligo.^21^

**Figure 4:**
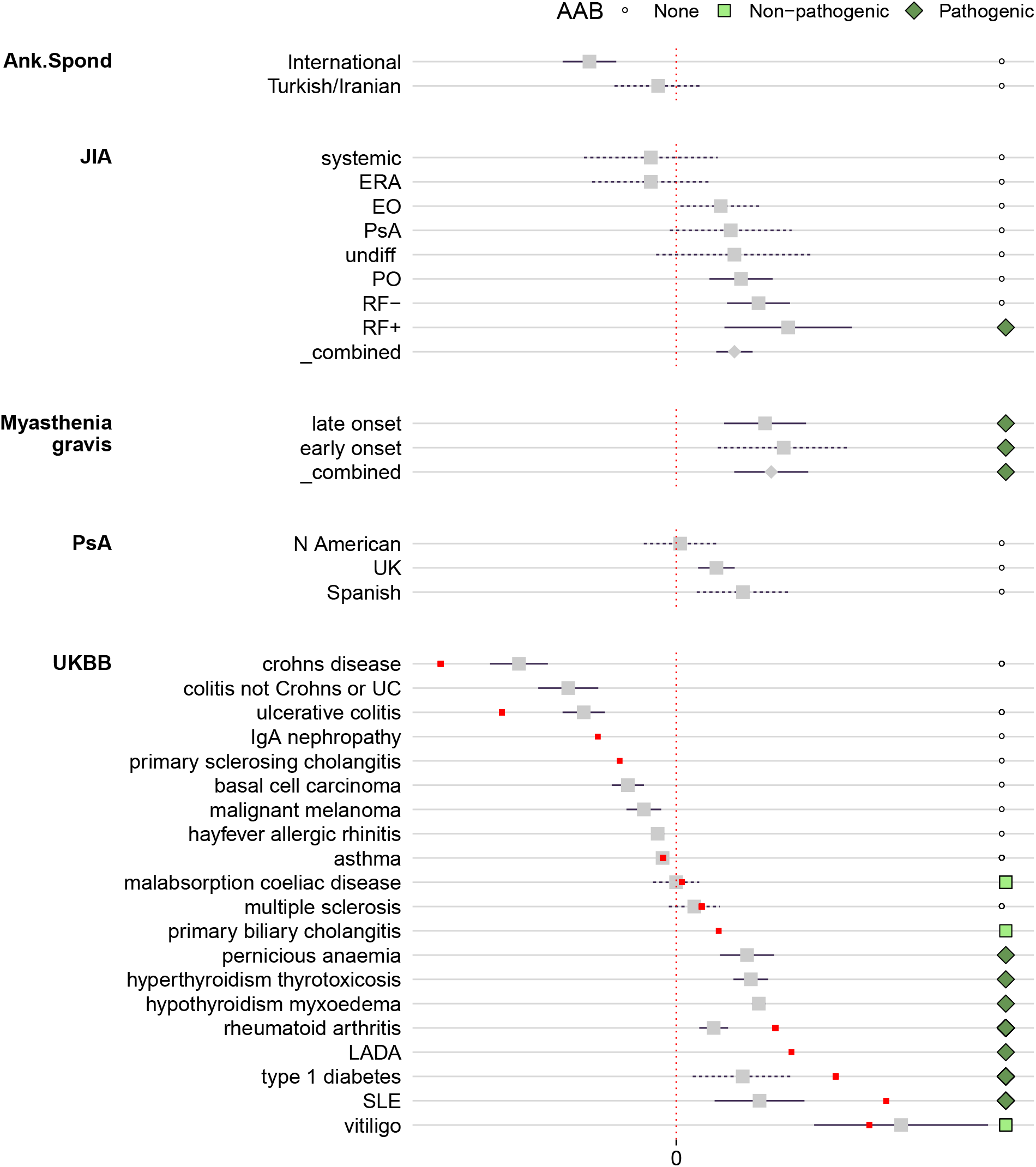
Forest plots showing projected values for diseases significant overall and on components 1. Grey squares dots indicate projected data and 95% confidence intervals. Red dots indicate the 13 IMD used for basis construction and for which no confidence interval is available. Points to the right of each line indicate disease classification according to whether they have specific autoantibodies that are either directly implicated in disease pathogenesis (“pathogenic”) or which are specific to the disease, but not involved in pathogenesis (“non-pathogenic”). Diseases that are not associated with specific autoantibodies were classified as “none”. Abbreviations: ANCA- = anti-neutrophil cytoplasmic antibody negative, Ank. Spond = ankylosing spondylitis, EGPA = eosinophilic granulomatosis with polyangiitis, EO = extended oligo, ERA = juvenile enthesitis-related arthritis, IgGPos = IgG positive, JIA = juvenile idiopathic arthritis, LADA = latent autoimmune diabetes in adults, NMO = neuromyelitis optica, PO = persistent oligo, PsA = psoriatic arthritis, RF +/- = polyoligo rheumatoid factor positive/negative, SLE = systemic lupus erythematosus, UC = ulcerative colitis.

To help characterise the biology captured by individual components we projected additional datasets: blood counts,^22^ immune cell counts^23^ and serum cytokine concentrations^24^ (Supplementary Tables 5, 6 and 7). Testing for consistency identified outliers in the blood count data, which had been generated from a much larger sample, and so we additionally filtered on consistency in that dataset. These data aided interpretation of two further components.

**PC13** was striking for the general association of many diseases across all four main clusters in a concordant direction, and was the only component for which any projected trait was more extreme than any original basis trait (Fig. 5). EGPA, which showed the most extreme projected values on this component of any diseases, is classified as an eosinophilic form of anti-neutrophil cytoplasmic antibody (ANCA)-associated vasculitis (AAV) and both asthma and raised eosinophil count are included in its diagnostic criteria. We found PC13 was strongly associated with higher eosinophil counts in a population cohort^22^ (FDR<10^-200^), suggesting that this component describes eosinophilic involvement in IMDs.

**Figure 5:**
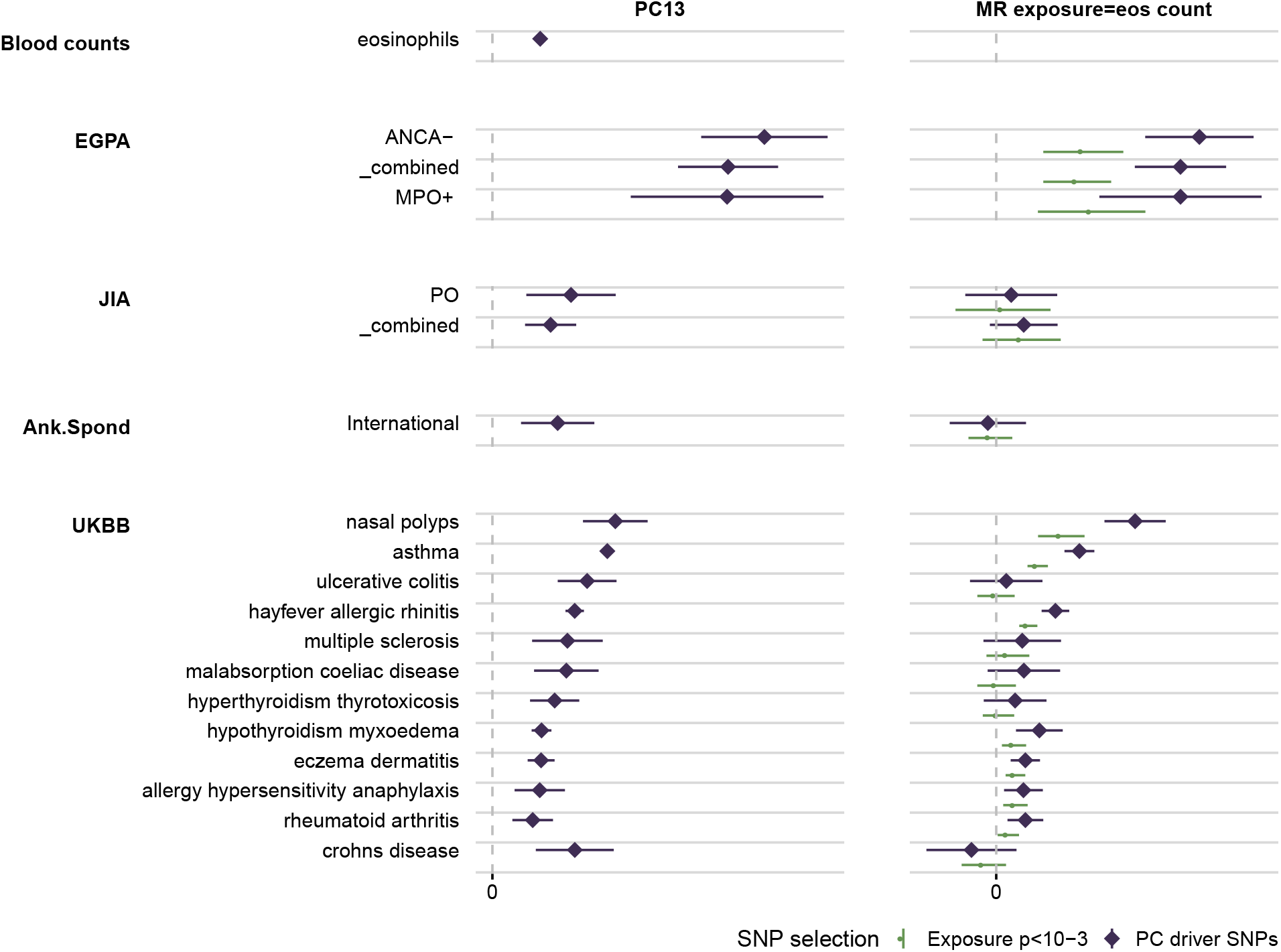
Forest plot of significant traits on PC13 (left panel) which also shows association with eosinophil counts in blood21. Mendelian randomisation (MR) analysis (right panel) using PC13 driver SNPs (purple) or SNPs significantly associated (*p* < 10^-3^) with eosinophil counts (green). Abbreviations: ANCA- = anti-neutrophil cytoplasmic antibody negative, Ank. Spond = ankylosing spondylitis, EGPA = eosinophilic granulomatosis with polyangiitis, JIA = juvenile idiopathic arthritis, MPO+ = myeloperoxidase-positive, PO = persistent oligo, RF- = polyoligo rheumatoid factor negative.

Eosinophils are pro-inflammatory leukocytes with an established role in atopic diseases such as asthma,^25^ inflammatory diseases such as IBD^26^ and autoimmune diseases such as RA.^27^ Mendelian Randomization (MR) analysis of blood cell traits had previously further associated eosinophils with celiac disease (CEL), asthma and T1D.^22^ We conducted MR analysis twice, first selecting SNPs according to significant association with eosinophil count and second using the driver SNPs for PC13. Results were similar, although estimates using PC13 driver SNPs tended to be larger, which suggests some heterogeneity, for example that only a subset of eosinophil-associated SNPs also associated with IMD risk. Our analysis thus confirms earlier findings, and extends the list of IMD with genetically supported involvement of eosinophils to include EGPA, JIA subtypes, AS, ATD, MS, hayfever and eczema, in agreement with other recent findings.^28^

**PC3** (Fig. 6) was the only component which showed a significant relationship with any serum cytokine concentration. Higher concentrations of CXCL9 (MIG) and CXCL10 (IP-10), Th1 chemoattractants and ligands to the regulator of leukocyte trafficking CXCR3, were both significant in the same direction as several autoimmune diseases, with strongest signals for myasthenia gravis, several JIA subtypes, as well as IBD, CEL, AS and sarcoidosis. In MR analysis, while PC3 driver SNPs predicted association of IP-10 and MIG with these IMDs, SNPs selected by significant association to cytokine levels themselves did not. This suggests that raised serum IP-10 and MIG are not themselves causally associated with IMD risk, but that these driver SNPs mark a risk factor that contributes to serum IP-10 and MIG.

**Figure 6:**
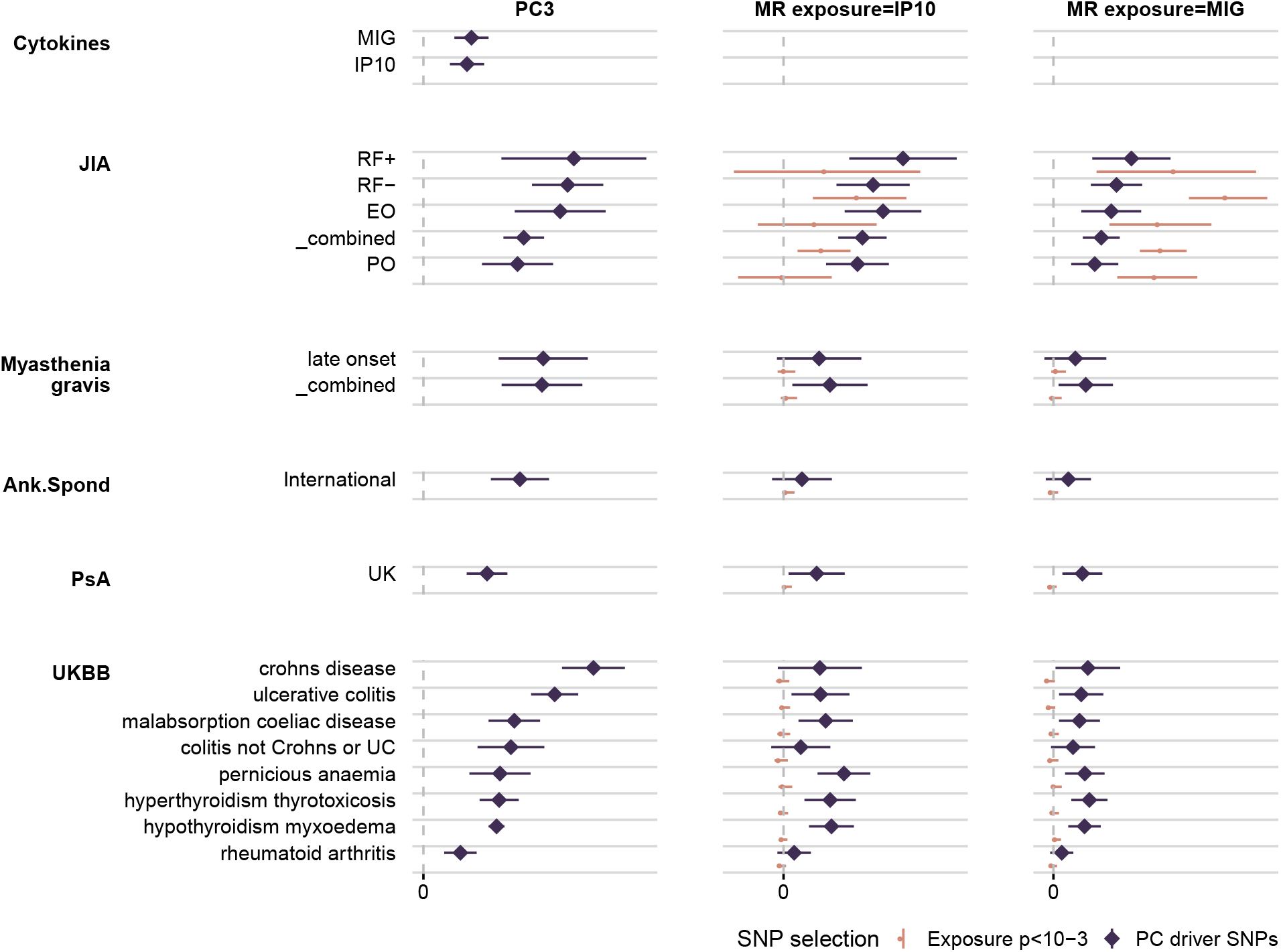
Forest plot of significant traits on PC3 (left panel) which also shows association with serum cytokine levels of IP-10 (CXCL10) and MIG (CXCL9). Mendelian randomisation analysis of IP-10 (middle panel) or MIG (right panel) using PC3 driver SNPs finds these cytokines to be associated with IMD risk (purple), but MR analysis using only SNPs significantly associated (*p* < 10^-3^) with either cytokine (green) do not show association. Abbreviations: EO = extended oligo, PO = persistent oligo, RF +/- = polyoligo rheumatoid factor positive/negative, UC = ulcerative colitis.

### Genetic distinctions within clinically heterogeneous and rare diseases

Our basis has only 13 dimensions. If the genetic susceptibility of rare IMD and IMD subtypes overlaps that of common IMD, we can increase power by focusing on these dimensions. Of 22 diseases or disease subtypes with < 1000 cases, 12 were significant (FDR<1%), even with as few as 132 cases (NMO IgGPos).

Most disease subtypes clustered together, even when not significant (Fig. 3). For example, myasthenia gravis, a chronic, autoimmune, neuromuscular disease characterized by muscle weakness, has been shown to have a bimodal incidence pattern by age, and some genetic associations have been identified only for the late onset subtype.^29^ However, both subtypes fell in very similar locations across all components, and cluster together along with several subtypes of JIA.

EGPA is a rare form of AAV (annual incidence 1-2 cases per million) for which genetic differences relating to autoantibody status have been identified.^11^ We included both myeloperoxidase (MPO) ANCA+ and ANCA-cases, as well as a study of non-eosinophilic MPO+ ANCA-associated vasculitis.^30^ While all forms of vasculitis fell on the adaptive immunity side on PC1, the EGPA subtypes typically resembled each other much more closely than the MPO+ EGPA resembled MPO+ ANCA-associated vasculitis, with EGPA showing a particularly strong signal on PC13, consistent with the diagnostic criteria which include overt eosinophilia.

For two other diseases, however, subtypes did not consistently cluster together. NMO is a rare (prevalence 0.03–0.4:10,000) disease affecting the optic nerve and spinal cord, for which HLA association is established^9^ and which can be divided according to aquaporin 4 autoantibody seropositivity status (IgG+ or IgG-). The projections of seropositive and seronegative NMO showed non-significant differences on several components, leading to differential clustering. While seropositive NMO clustered with the classical autoimmune diseases, most closely with SLE and Sjögren’s disease, IgG- NMO clustered away from the classic seropositive diseases, most closely with MS. This finding mirrors analysis which directly compared NMO subtypes to each of SLE and MS via polygenic scores,^9^ and strengthens the findings by specifically identifying SLE and MS as the nearest neighbours of IgG+ and IgG- NMO respectively, out of 60 IMDs considered for clustering.

JIA is a heterogeneous paediatric disease, with an overall childhood prevalence in Europe of 20:10,000,^31^ and with seven recognised subtypes.^32^ While studies have begun to identify distinct genetics of the systemic subtype^33^ and have shown subtype-specific differences in the MHC,^34^ systematic comparison between subtypes has been underpowered. Although, the systemic and enthesitis-related arthritis (ERA) subtypes were not significant despite relatively moderate sample sizes (219 and 267 cases respectively), they clustered together with MS and AS respectively, apart from the other JIA subtypes which clustered with the other autoimmune diseases.

### Mapping component-level associations to SNPs

Given that most of the IMD and subtypes with small GWAS studies have few established genetic associations, we sought to exploit the component-level associations above to detect new disease associations. We found a strong enrichment for small GWAS p values at driver SNPs on trait-significant components (Supplementary Fig. 19). Using a “subset-selected” FDR approach,^35^ we analysed driver SNPs for 22 significant trait-component pairs (12 unique traits), and identified 25 trait-SNP associations (subset-selected FDR < 1%, Table 1) after pruning SNPs in LD. Twelve of these were genome-wide significant (p < 5×10^-8^) either in this study (4 associations) or in other published data (8 associations) and a further five were significant in other published analysis that levered external data. These included, for example, the non-synonymous *PTPN22* SNP rs2476601 which was associated with myasthenia gravis (overall and the late onset subset) by subset-selected FDR < 0.01. This SNP was previously associated with myasthenia gravis in a different study,^36^ and lack of clear replication in the data analysed here (SNP P=6×10^-5^) was attributed to differences in population structure. Eight associations (five variants) were not previously reported to our knowledge, including associations near *IRF1/IL5* for myasthenia gravis, near *TNFSF11* for rheumatoid factor negative (RF-) JIA and near *CD2/CD2* 8 for EGPA.

**Table 1:**
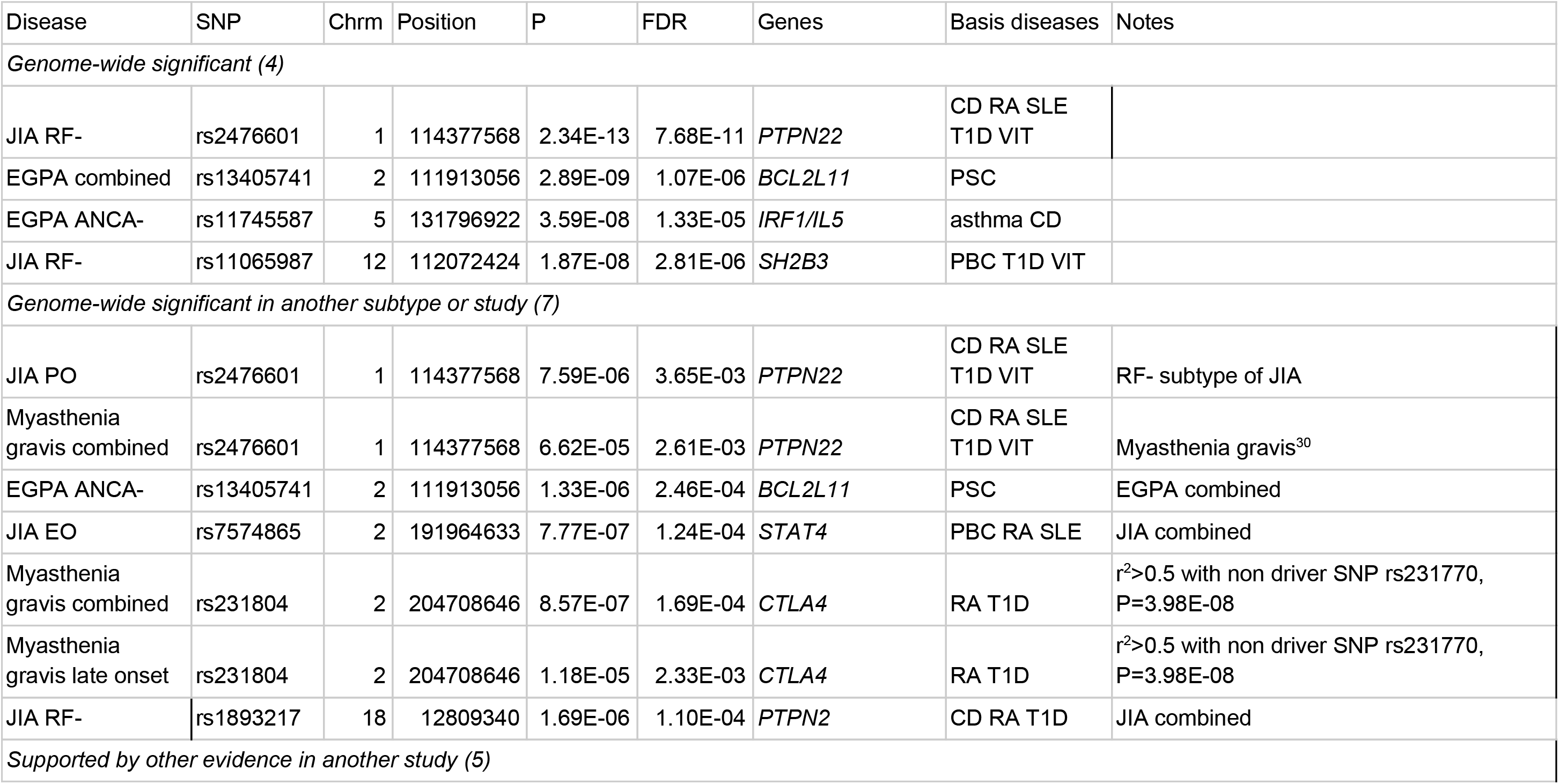
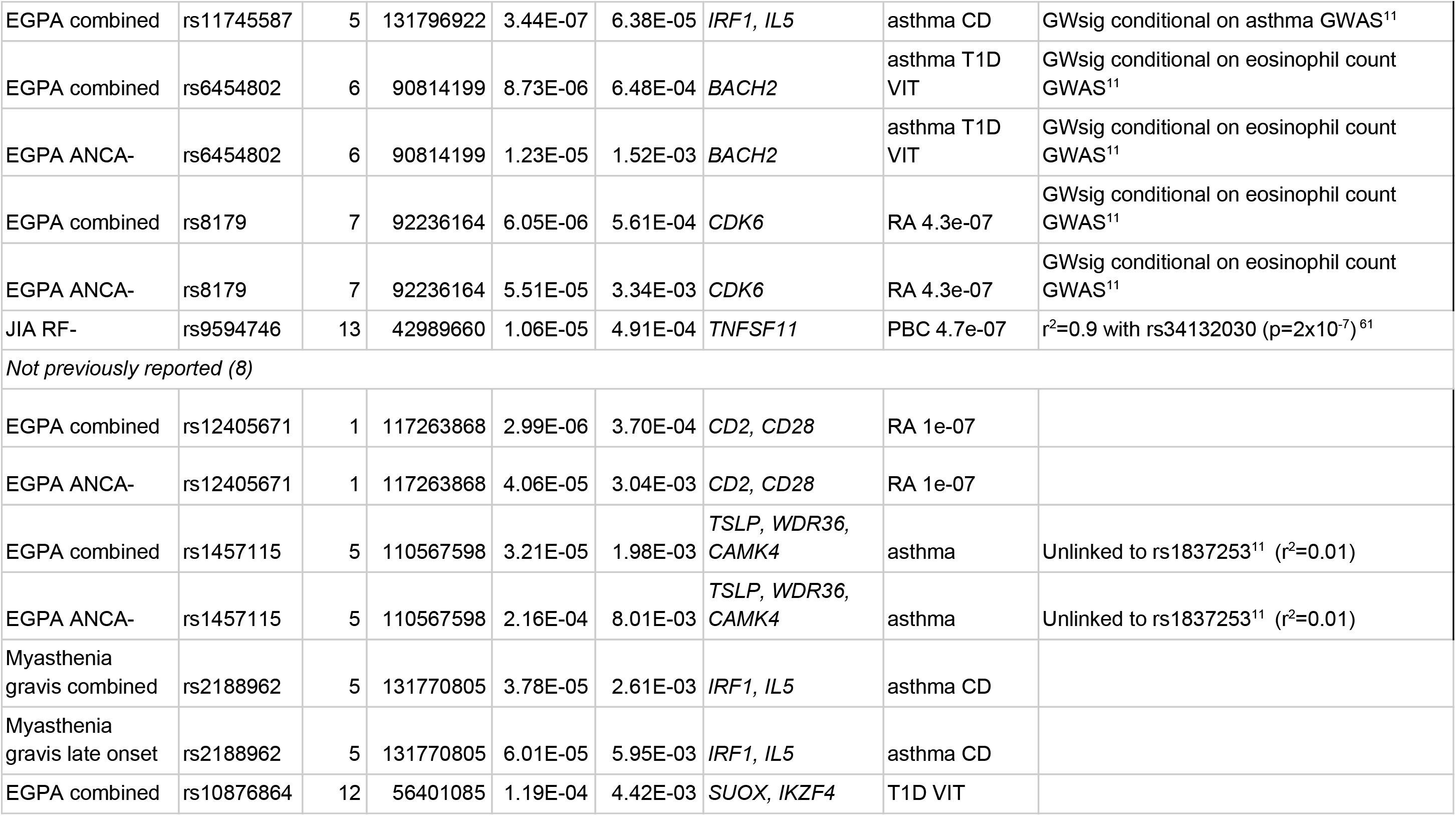
Disease-associated SNPs identified through subset-selected FDR (ssFDR) < 0.01 amongst driver SNPs belonging to disease-significant components. Genes listed are nearby genes previously mentioned in the literature for the listed disease or basis diseases associated to this SNP, and are intended to indicate location; no evidence for gene causality has been assessed here. Where no basis diseases are associated with the SNP at genome-wide significant threshold (GWsig, p<5×10^-8^), the strongest association and its p value are shown.

## Discussion

Our motivation in this work was threefold. First, to overcome the problems of dimensionality to allow an overview of genetic association patterns from multiple related diseases without over-simplification. While previous efforts to relate different traits through GWAS statistics have focused on large studies and shown that they can distinguish broad classes of IMD, cardiovascular and metabolic diseases,^5,13^ we have tackled the problem of finding structure *within* a single class of diseases. Unlike other applications of PCA to genetics, we split our datasets into “training” and “test” sets, enabling standard statistical hypothesis testing and providing robustness against overfitting.

Our second motivation was to extract different axes underlying IMD genetic risk. Work in metabolic^37^ and psychiatric^38^ diseases have taken related approaches to attempt to learn composite factors underlying risk of these related diseases through deeper phenotyping of patients before testing these factors for genetic association. Alternatively, decomposition of estimated effects at 94 type 2 diabetes risk variants, together with their effects on 46 metabolic traits was used to cluster these variants into 5 groups, three focused on insulin resistance and two on beta cell function.^39^ Here, we hoped to learn the same sorts of factors by decomposing only summary GWAS data on clinical disease endpoints. Our continuous shrinkage weight learnt across all training datasets enables us to extract disease-relevant structure, with projected traits lying close to their training data counterparts, something achieved with disease-specific hard thresholded weights^14^ for only the largest datasets.

We indeed found variable contribution of different components to different disease subsets. The autoimmune/(auto-)inflammatory axis in IMD represented by PC1 is well documented, with the gradient along PC1 corresponding to a shift from auto-antibody seronegative to seropositive diseases. Significant IMD on the MIG/IP-10-associated PC3 included both ‘seropositive’ and ‘seronegative’ diseases, although not atopy, while all three groups were represented on the eosinophil associated PC13. IP-10 and MIG are chemokines, secreted by epithelial and dendritic cells (amongst others), which act as chemoattractants for immune cells which express the receptor CXCR3, including Th1 cells. Both MIG and IP-10 expression at the site of autoimmune target have been implicated in the development of autoimmunity^40,41^ and IP-10 has been observed to be upregulated in follicular cells of patients with myasthenia gravis.^42^ Serum IP-10 has also been found to be raised in patients with recent onset T1D,^43,44^ and Graves’ disease (hyperthyroidism)^41^ and to correlate with increased disease activity in SLE^45^ and AS.^46^ While these observations support a link between certain IMD and serum cytokine levels, our results do not directly implicate these cytokines as causal. Both cytokines and blood count data were measured in unselected population cohorts which will include individuals with IMD, such that the association with IMD may be causal or consequential. We suggest that PC3 represents an IMD-related process that contributes to serum cytokine levels. Nonetheless, clinical efficacy of MDX1100, a monoclonal antibody to IP-10, has been demonstrated in RA^47^ and a dose-response relationship observed in UC^48^ and our results suggest IP-10 blockade might also be considered in patients with myasthenia gravis, JIA, AS, CEL and sarcoidosis.

Our final motivation was to exploit the lower dimensional representation to generate new knowledge in rare IMDs. The number of polymorphic human genetic variants together with our understanding that genetic effects on human disease are generally modest has lead to massive GWAS to overcome the penalty that must be applied for multiple testing. This is simply not possible for rare diseases. One of the tools which has enhanced rare disease GWAS is the borrowing of information from larger GWAS of aetiologically related diseases^11^ and our basis serves a similar function here, by leveraging information about a SNP’s potential to be IMD-associated, we can both increase genetic discovery and place less common diseases in the context of their more prevalent counterparts. More generally, studies of SRD are being enabled on massive scale by UKBB^49^ and 23andMe,^50^ although studies of such cohorts tend to focus on the more common diseases such as type 2 diabetes and coronary heart disease. Our results provide reassurance that SRD associations are consistent with those from targeted GWAS studies, and extend their utility to IMD and other diseases which are generally found at a lower frequency.

Our approach represents genetic associations for aetiologically related traits in a reduced dimension basis, with attached estimates of uncertainty. This will enable novel cross-diseases analyses, and thus increase understanding of both the underlying components of disease risk as well as placing lower prevalence diseases in context of their related common diseases.

## Methods

### Summary statistic datasets

We constructed a compendium of publicly available GWAS summary statistics across a wide range of traits including UKBB traits (http://www.nealelab.is/uk-biobank, http://geneatlas.roslin.ed.ac.uk/ - Supplementary Tables 2 and 4), IMD relevant GWAS studies (Supplementary Table 3), and GWAS of quantitative measures from blood count data^22^, immune cell counts^23^ and cytokine levels^24^ (Supplementary tables 5, 6 and 7).

Disease GWAS data were obtained from the URL given in Supplementary Tables 1, 3, or via request to study authors, with the exception of those listed below.

### Vasculitis GWAS analysis

AAV belongs to a group of IMD characterised by inflammation of the small and medium-sized blood vessels with evidence of circulating pathogenic autoantibodies. It comprises three main syndromes: granulomatosis with polyangiitis (GPA), microscopic polyangiitis (MPA) and EGPA. The two primary antigenic targets of ANCA are proteinase 3 (PR3) and myeloperoxidase (MPO). Although PR3-ANCA is the predominant serotype in GPA and MPO-ANCA is more commonly found in MPA, there is a significant overlap between these syndromes.

The vasculitis cohort used to construct the basis was part of the discovery cohort from the AAV GWAS performed by the European Vasculitis Genetics Consortium,^30^ comprising 478 PR3-AAV cases, 264 MPO-AAV and 5,259 controls from the Wellcome Trust Case Control Consortium. All cases had a clinical diagnosis of either GPA or MPA according to the European Medicines Agency algorithm, supported by a positive ANCA assay. The genotyping, calling and data QC have been previously described.^30^. Briefly, the genotyping was performed by AROS Applied Biotechnology (Arthus, Denmark) using the Affymetrix SNP6 platform. Pre-phasing and genome-wide SNP imputation were performed using Eagle2 and Minimac3 respectively on the Michigan Imputation Server v1.0.3 that facilitates access to the HRC reference panel (HRC version r1.1 2016).^51^ Post-imputation, SNPs with MAF < 0.01 or r^2^ < 0.3 were removed from dataset using BCFtools version 1.2, leaving a total of 7,656,576 SNPs available for case-control association testing using a linear mixed model with BOLT-LMM software v2.3.2.^52,53^

### JIA and PsA GWAS analysis

The JIA and PsA GWAS datasets were generated and QC’d using the same strategies.

#### Genotyping and statistical quality control

JIA and PsA DNA samples were genotyped on the Illumina Infinium CoreExome genotyping array in accordance to the manufacturer’s instructions at the Centre for Genetics and Genomics Versus Arthritis (The University of Manchester). Genotype calling was performed by the GenCall algorithm in the GenomeStudio Data Analysis software platform (Genotyping Module v1.8.4). Preliminary genotype clustering was performed using the default Illumina cluster file to identify poor quality samples (call rate < 0.90). Following exclusion of low quality samples automated reclustering was performed to calibrate genotype clusters based on the study samples. Sample-level quality control (QC) was performed based on the following exclusion criteria: final call rate < 0.98, outlier based on autosomal heterozygosity (2 standard deviations from the mean) and discrepancy between genetically inferred sex and database records. SNPs were excluded if they were non-autosomal, call rate < 0.98 or a minor allele frequency < 0.01. PsA was compared with 4596 controls from the WTCCC2 study (REF). JIA was compared with 9,965 population controls from the UK Household Longitudinal Study (https://www.understandingsociety.ac.uk/) accessed via the European Genotype-phenome Archive. Samples were genotyped at the Wellcome Trust Sanger Institute using the Illumina Infinium CoreExome genotyping array. Sample and SNP QC is consistent with that described above for case samples.

Case and control datasets were combined retaining the intersection of SNPs. Identity-by-descent was used to identify related individuals (kinship coefficient > 0.0884) across all study samples performed with the KING software package (version 1.9). For each related pair the sample with the highest call rate was preferentially retained. Individuals were excluded if they were identified as outliers based on ancestry using principal component analysis (PCA) performed with the flashpca software package (version 2.0) where outliers were identified using aberrant R library (version 1.0).

#### Imputation

Prior to imputation SNPs with ambiguous alleles (C/G and A/T) were excluded and remaining SNPs were aligned to the Haplotype Reference Consortium (HRC) panel (version 1.1) using the HRC imputation preparation tool (https://www.well.ox.ac.uk/~wrayner/tools/). Imputation was performed using the Michigan Imputation server where phasing was performed with Shapeit2 and the HRC panel. Following imputation SNPs were excluded based on a MAF < 0.01 and imputation accuracy (r^2^) < 0.4.

#### Association testing, PsA

case-control association testing was performed using the SNPTEST software package (version 2.5.2) using the score method to account for imputation uncertainty. Three principal components, calculated as described above, were included as covariates to account for any residual population structure.

#### Association testing, JIA

case-control association testing was performed using the snp.rhs.estimates function in the R package SnpStats, comparing in turn overall, or JIA subtypes to the control group. Three principal components, calculated as described above, were included as covariates to account for any residual population structure.

### Construction of basis

There are three particular challenges with performing PCA on GWAS summary statistics. First, the SNP effect estimates must be on the same scale; second, we must deal with variable correlation between input dimensions (SNPs) due to LD; and third, while all SNPs are expected to show small deviations between studies due to random noise, different genotyping platforms and data processing decisions, only a minority of SNPs will be truly related to the diseases of interest.

The uncertainty attached to 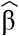 depends on both study sample size and SNP minor allele frequency (MAF). We adjusted for the variance due to MAF, σ^2^_*MAF*_, as this varies between SNPs, but not variance due to sample size, as this would overly shrink smaller studies relative to larger. We dealt with the second two challenges simultaneously, using a Bayesian fine mapping technique which calculates the posterior probability that each SNP is causal for each trait, under the assumption that at most one causal variant exists in each recombination hotspot-defined block of SNPs ^54,55^. At each SNP, we computed a weighted average of the posterior probabilities across input studies to create an overall weight for that SNP, *w* . *w* will be close to zero when there is no association in a region, limiting the effects of technical noise between studies, and will otherwise act to weight associated SNPs according to the extent of LD in a region. The final input for basis creation is a matrix of 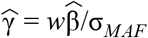.

We identified 13 IMD GWAS studies with >6,000 samples of European ancestry for which full summary statistics were publicly available (Supplementary Table 1). We selected SNPs present in all 13 studies, with MAF>1% in the 1000 Genomes Phase 3 EUR data. Additionally we excluded SNPs overlapping the MHC region (GrCh37 Chr6:20-40Mb) or for which the unambiguous assignment of the effect allele was impossible (e.g. palindromic SNPs). We harmonised all effect estimates to be with respect to the alternative allele relative to the reference allele as defined by the 1000 genome reference genotype panel. After filtering, harmonised effect estimates were available for 265,887 SNPs across all 13 selected ‘basis’ traits. In order to provide a baseline for subsequent analyses we created an additional synthetic ‘control’ trait, for which effect sizes across all traits were set to zero. We used these to construct two matrices *M* and *M’* where elements reflect raw 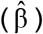 and shrunk effect sizes 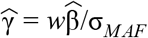 respectively, such that, rows and columns reflect traits (n=14) and SNPs (p=265,887). After mean centring columns we used the R command *prcomp* to carry out PCA of both *M* and *M’* to generate naive and “shrunk” IMD bases, retaining *m=n*-1=13 components, which corresponded to the fewest components needed to minimise the mean squared reconstruction error (Supplementary Fig 20).

We noted that the majority of entries in the *p* x *m* PCA rotation matrix, *R*, were close to 0, and chose to hard threshold these to 0 for computational efficiency and to identify which *driver SNPs* were relevant to each component. To do this, using *R_k_* to represent the *k*^th^ column of *R*, we define R_k_(*α*)=R_k_ x I(|R_k_|>*α*) where I() is an indicator function and “x” represents element-wise multiplication. We quantify the distance between projection with *R_k_* and *R_k_*(*α*) by

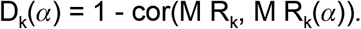

We chose the threshold for each component, *α*_k_, as the largest value *α* such that D_k_(*α*) < 0.001. Finally, we defined the sparse basis rotation matrix Q as the matrix constructed from the column vectors R_k_, k=1,…,m. This identified both driver SNPs which define the support for each component, and enabled computationally efficient examination of many traits in the reduced dimension space defined.

### Projection

Prior to projection, effect alleles were aligned to the 1000 genome reference genotype panel. For traits sensitive to missing data (studies of NMO^9^ and PsA by Aterido,^56^ see Supplementary Note), we imputed missing variants using ssimp^57^ (v 0.5.6 ----ref 1KG/EUR --impute.maf 0.01), otherwise we set effect estimates to zero. Data were then shrunk as for the basis traits (multiplying by *w*/σ_*MAF*_), and projected into basis space by multiplying by the sparse basis rotation matrix Q. The locations of all basis and projected traits in this space are reported as 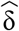, ie relative to the location of a synthetic “control” trait vector of **0** also projected into the space. We calculated variance of the projected values (Supplementary Note) and quantified consistency using a weighted Spearman rank correlation on a subset of driver SNPs in low LD (r^2^<0.01), with weights *w*/σ_*MAF*_ and significance determined by permuting the projected values (Supplementary Note). All projected values are given in Supplementary Table 8.

We used the *hclust()* function in R to cluster diseases in the basis using agglomerative hierarchical clustering according to Ward’s criterion (method=“Ward.D2”) on the Euclidean distance between projected locations of each disease in the basis.

We calculated p values for null hypotheses that the vector δ = 0 across all 13 components using a chisq test (Supplementary Note). We called significant associations according to FDR < 0.01, calculated using the Benjamini-Hochberg approach, and run independently within the broad categories: primary analysis (UKBB self reported disease and cancer, plus IMD-relevant GWAS); blood cell counts; cytokines; immune cell counts. This was our primary measure of significance. We took the same strategy to independently calculate FDR for each component individually for additional annotation, and traits were considered “component-significant” if they were significant on that component *and* overall.

Classification of diseases according to autoantibody status was performed by a specialist clinician using available medical literature. This assignment was blinded to the PC1 results.

### Proportionality of effects across different datasets for the same trait

We tested the null hypothesis of proportionality using the coloc.test function in the coloc package^58^ which takes into account the uncertainty in the projection estimate, assuming different PCs are independent. Small p values in this test correspond to the observed data being unlikely under the null of proportionality, and would suggest that studies of the same disease in different populations were not comparable.

### Candidate significant driver SNPs

For each of 10 diseases or subtypes with < 2000 cases and significant on at least one component (Myasthenia gravis, late onset; EGPA, MPO+, ANCA- and combined; JIA, extended oligo (EO), persistent oligo (PO), RF-, and RF+), we selected all driver SNPs on any significant component, and calculated the FDR within this set of SNPs as a subset-selected FDR.^35^ We ordered SNPs by increasing values of ssFDR, and deleted any SNPs in the list that were in LD (r^2^>0.1) with a higher placed SNP, leaving a set of unlinked SNPs associated with each trait shown in Table 1. These were annotated through literature searches.

### Code availability

An R implementation of the method is available from https://github.com/ollyburren/cupcake. Code to run the analyses presented here is available from https://github.com/ollyburren/imd-basis. Code underlying the online tool to allow projection of additional data into the basis (https://grealesm.shinyapps.io/IMDbasisApp/) is available at https://github.com/GRealesM/IMDbasisApp.

## Author contributions

Conceived the study, drafted paper: CW, OB.

Wrote the software: OB.

Performed analyses: OB, CW, LW, JB.

Interpreted data: OB, CW, JL, PDWK, KGCS.

Acquired data: OB, JB, WT, JB, KGCS, PAL, AB, CW.

Created online projection tool: GR, OB.

All authors read and approved the manuscript.

## Acknowledgements

We thank the following for sharing data from their studies:

Ann Morgan and Jennifer Barrett for methotrexate response in RA, on behalf of the CARDERA, IACON, PAMERA, and RAMS Consortia;^59^

Jonas Kuiper and Bobby Koeleman for the birdshot chorioretinopathy GWAS;^60^

We thank Urs Christen for helpful discussions on IP-10.

This study acknowledges the use of the following UK JIA cohort collections: British Society of Paediatric and Adolescent Rheumatology (BSPAR) study group, Childhood Arthritis Prospective Study (CAPS) (funded by Versus Arthritis, grant reference number 20542), Childhood Arthritis Response to Medication Study (CHARMS) (funded by Sparks UK, reference 08ICH09, and the Medical Research Council, reference MR/M004600/1), United Kingdom Juvenile Idiopathic Arthritis Genetics Consortium (UKJIAGC). Genotyping of the UK JIA and PsA case samples was supported by the Versus Arthritis grants reference numbers 20385 and 21754. This research was funded by the NIHR Manchester Biomedical Research Centre and supported by the Manchester Academic Health Sciences Centre (MAHSC). The views expressed are those of the author(s) and not necessarily those of the NHS, the NIHR or the Department of Health. We would like to acknowledge the assistance given by IT Services and the use of the Computational Shared Facility at The University of Manchester. Understanding Society: The UK Household Longitudinal Study is led by the Institute for Social and Economic Research at the University of Essex and funded by the Economic and Social Research Council. The survey was conducted by NatCen and the genome-wide scan data were analysed and deposited by the Wellcome Trust Sanger Institute. Information on how to access the data can be found on the Understanding Society website https://www.understandingsociety.ac.uk/.

## References

1. Buniello, A. et al. The NHGRI-EBI GWAS Catalog of published genome-wide association studies, targeted arrays and summary statistics 2019. Nucleic Acids Res. 47, D1005–D1012 (2019).

2. Cotsapas, C. & Hafler, D. A. Immune-mediated disease genetics: the shared basis of pathogenesis. Trends Immunol. 34, 22–26 (2013).

3. Majumdar, A., Haldar, T., Bhattacharya, S. & Witte, J. S. An efficient Bayesian meta-analysis approach for studying cross-phenotype genetic associations. PLoS Genet. 14, (2018).

4. Cotsapas, C. et al. Pervasive sharing of genetic effects in autoimmune disease. PLoS Genet. 7, e1002254 (2011).

5. Bulik-Sullivan, B. et al. An atlas of genetic correlations across human diseases and traits. Nat. Genet. 47, 1236–1241 (2015).

6. Fortune, M. D. et al. Statistical colocalization of genetic risk variants for related autoimmune diseases in the context of common controls. Nat. Genet. 47, 839+ (2015).

7. Yang, Y. et al. Clinical whole-exome sequencing for the diagnosis of mendelian disorders. N. Engl. J. Med. 369, 1502–1511 (2013).

8. Ouwehand, W. H. Whole-genome sequencing of rare disease patients in a national healthcare system. bioRxiv 507244 (2019) doi:10.1101/507244.

9. Estrada, K. et al. A whole-genome sequence study identifies genetic risk factors for neuromyelitis optica. Nat. Commun. 9, 1929 (2018).

10. Li, J. et al. Association of CLEC16A with human common variable immunodeficiency disorder and role in murine B cells. Nat. Commun. 6, 6804 (2015).

11. Lyons, P. A. et al. Genome-wide association study of eosinophilic granulomatosis with polyangiitis reveals genomic loci stratified by ANCA status. Nat. Commun. 10, 5120 (2019).

12. Price, A. L. et al. Principal components analysis corrects for stratification in genome-wide association studies. Nat. Genet. 38, 904–909 (2006).

13. Chang, D. & Keinan, A. Principal component analysis characterizes shared pathogenetics from genome-wide association studies. PLoS Comput. Biol. 10, e1003820 (2014).

14. Tanigawa, Y. et al. Components of genetic associations across 2,138 phenotypes in the UK Biobank highlight adipocyte biology. Nat. Commun. 10, 4064 (2019).

15. Sudlow, C. et al. UK biobank: an open access resource for identifying the causes of a wide range of complex diseases of middle and old age. PLoS Med. 12, e1001779 (2015).

16. McGonagle, D. & McDermott, M. F. A proposed classification of the immunological diseases. PLoS Med. 3, e297 (2006).

17. Boniface, K., Seneschal, J., Picardo, M. & Taïeb, A. Vitiligo: Focus on Clinical Aspects, Immunopathogenesis, and Therapy. Clin. Rev. Allergy Immunol. 54, 52–67 (2018).

18. Yuan, Y. et al. Identification of Novel Autoantibodies Associated With Psoriatic Arthritis. Arthritis Rheumatol 71, 941–951 (2019).

19. Singh, H., Nugent, Z., Demers, A. A. & Bernstein, C. N. Increased risk of nonmelanoma skin cancers among individuals with inflammatory bowel disease. Gastroenterology 141, 1612–1620 (2011).

20. Singh, S. et al. Inflammatory bowel disease is associated with an increased risk of melanoma: a systematic review and meta-analysis. Clin. Gastroenterol. Hepatol. 12, 210–218 (2014).

21. Toh, B.-H. Pathophysiology and laboratory diagnosis of pernicious anemia. Immunol. Res. 65, 326–330 (2017).

22. Astle, W. J. et al. The Allelic Landscape of Human Blood Cell Trait Variation and Links to Common Complex Disease. Cell 167, 1415–1429.e19 (2016).

23. Roederer, M. et al. The genetic architecture of the human immune system: a bioresource for autoimmunity and disease pathogenesis. Cell 161, 387–403 (2015).

24. Ahola-Olli, A. V. et al. Genome-wide Association Study Identifies 27 Loci Influencing Concentrations of Circulating Cytokines and Growth Factors. Am. J. Hum. Genet. 100, 40–50 (2017).

25. Busse, W. W. & Sedgwick, J. B. Eosinophils in asthma. Ann. Allergy 68, 286–290 (1992).

26. Al-Haddad, S. & Riddell, R. H. The role of eosinophils in inflammatory bowel disease. Gut vol. 54 1674–1675 (2005).

27. Hällgren, R., Feltelius, N., Svenson, K. & Venge, P. Eosinophil involvement in rheumatoid arthritis as reflected by elevated serum levels of eosinophil cationic protein. Clin. Exp. Immunol. 59, 539–546 (1985).

28. Diny, N. L., Rose, N. R. & Čiháková, D. Eosinophils in Autoimmune Diseases. Front. Immunol. 8, 484 (2017).

29. Renton, A. E. et al. A genome-wide association study of myasthenia gravis. JAMA Neurol. 72, 396–404 (2015).

30. Lyons, P. A. et al. Genetically distinct subsets within ANCA-associated vasculitis. N. Engl. J. Med. 367, 214–223 (2012).

31. Thierry, S., Fautrel, B., Lemelle, I. & Guillemin, F. Prevalence and incidence of juvenile idiopathic arthritis: a systematic review. Joint Bone Spine 81, 112–117 (2014).

32. Petty, R. E. et al. International League of Associations for Rheumatology classification of juvenile idiopathic arthritis: second revision, Edmonton, 2001. J. Rheumatol. 31, 390–392 (2004).

33. Ombrello, M. J. et al. Genetic architecture distinguishes systemic juvenile idiopathic arthritis from other forms of juvenile idiopathic arthritis: clinical and therapeutic implications. Ann. Rheum. Dis. 76, 906–913 (2017).

34. Hinks, A. et al. Fine-mapping the MHC locus in juvenile idiopathic arthritis (JIA) reveals genetic heterogeneity corresponding to distinct adult inflammatory arthritic diseases. Ann. Rheum. Dis. 76, 765–772 (2017).

35. Yekutieli, D. et al. Approaches to multiplicity issues in complex research in microarray analysis. Stat. Neerl. 60, 414–437 (2006).

36. Gregersen, P. K. et al. Risk for myasthenia gravis maps to a (151) Pro→Ala change in TNIP1 and to human leukocyte antigen-B*08. Ann. Neurol. 72, 927–935 (2012).

37. Avery, C. L. et al. A phenomics-based strategy identifies loci on APOC1, BRAP, and PLCG1 associated with metabolic syndrome phenotype domains. PLoS Genet. 7, e1002322 (2011).

38. Mallard, T. T. et al. Not just one p: Multivariate GWAS of psychiatric disorders and their cardinal symptoms reveal two dimensions of cross-cutting genetic liabilities. bioRxiv 603134 (2019) doi:10.1101/603134.

39. Udler, M. S. et al. Type 2 diabetes genetic loci informed by multi-trait associations point to disease mechanisms and subtypes: A soft clustering analysis. PLoS Med. 15, e1002654 (2018).

40. Christen, U., McGavern, D. B., Luster, A. D., von Herrath, M. G. & Oldstone, M. B. A. Among CXCR3 chemokines, IFN-gamma-inducible protein of 10 kDa (CXC chemokine ligand (CXCL) 10) but not monokine induced by IFN-gamma (CXCL9) imprints a pattern for the subsequent development of autoimmune disease. J. Immunol. 171, 6838–6845 (2003).

41. Romagnani, P. et al. Expression of IP-10/CXCL10 and MIG/CXCL9 in the thyroid and increased levels of IP-10/CXCL10 in the serum of patients with recent-onset Graves’ disease. Am. J. Pathol. 161, 195–206 (2002).

42. Meraouna, A. et al. The chemokine CXCL13 is a key molecule in autoimmune myasthenia gravis. Blood 108, 432–440 (2006).

43. Shimada, A. et al. Elevated serum IP-10 levels observed in type 1 diabetes. Diabetes Care 24, 510–515 (2001).

44. Antonelli, A. et al. Serum Th1 (CXCL10) and Th2 (CCL2) chemokine levels in children with newly diagnosed Type 1 diabetes: a longitudinal study. Diabet. Med. 25, 1349–1353 (2008).

45. Kong, K. O. et al. Enhanced expression of interferon-inducible protein-10 correlates with disease activity and clinical manifestations in systemic lupus erythematosus. Clin. Exp. Immunol. 156, 134–140 (2009).

46. Wang, J. et al. Circulating levels of Th1 and Th2 chemokines in patients with ankylosing spondylitis. Cytokine 81, 10–14 (2016).

47. Yellin, M. et al. A phase II, randomized, double-blind, placebo-controlled study evaluating the efficacy and safety of MDX-1100, a fully human anti-CXCL10 monoclonal antibody, in combination with methotrexate in patients with rheumatoid arthritis. Arthritis Rheum. 64, 1730–1739 (2012).

48. Mayer, L. et al. Anti-IP-10 antibody (BMS-936557) for ulcerative colitis: a phase II randomised study. Gut 63, 442–450 (2014).

49. Bycroft, C. et al. The UK Biobank resource with deep phenotyping and genomic data. Nature 562, 203–209 (2018).

50. Tian, C. et al. Genome-wide association and HLA region fine-mapping studies identify susceptibility loci for multiple common infections. Nat. Commun. 8, 599 (2017).

51. Das, S. et al. Next-generation genotype imputation service and methods. Nat. Genet. 48, 1284–1287 (2016).

52. Loh, P.-R. et al. Contrasting genetic architectures of schizophrenia and other complex diseases using fast variance-components analysis. Nat. Genet. 47, 1385–1392 (2015).

53. Loh, P.-R., Kichaev, G., Gazal, S., Schoech, A. P. & Price, A. L. Mixed-model association for biobank-scale datasets. Nat. Genet. 50, 906–908 (2018).

54. Wakefield, J. Bayes factors for genome-wide association studies: comparison with P-values. Genet. Epidemiol. 33, 79–86 (2009).

55. The Wellcome Trust Case Control Consortium et al. Bayesian refinement of association signals for 14 loci in 3 common diseases. Nat. Genet. 44, 1294–1301 (2012).

56. Aterido, A. et al. Genetic variation at the glycosaminoglycan metabolism pathway contributes to the risk of psoriatic arthritis but not psoriasis. Ann. Rheum. Dis. 78, (2019).

57. Rüeger, S., McDaid, A. & Kutalik, Z. Evaluation and application of summary statistic imputation to discover new height-associated loci. PLoS Genet. 14, e1007371 (2018).

58. Wallace, C. Statistical testing of shared genetic control for potentially related traits. Genet. Epidemiol. 37, 802–813 (2013).

59. Taylor, J. C. et al. Genome-wide association study of response to methotrexate in early rheumatoid arthritis patients. Pharmacogenomics J. 18, 528–538 (2018).

60. Kuiper, J. J. W. et al. A genome-wide association study identifies a functional ERAP2 haplotype associated with birdshot chorioretinopathy. Hum. Mol. Genet. 23, 6081–6087 (2014).

61. Hinks, A. et al. Dense genotyping of immune-related disease regions identifies 14 new susceptibility loci for juvenile idiopathic arthritis. Nature Publishing Group 45, 664–669 (2013).

