## Supplementary Figures for "Informed dimension reduction of clinically-related genome-wide association summary data characterises cross-trait axes of genetic risk"

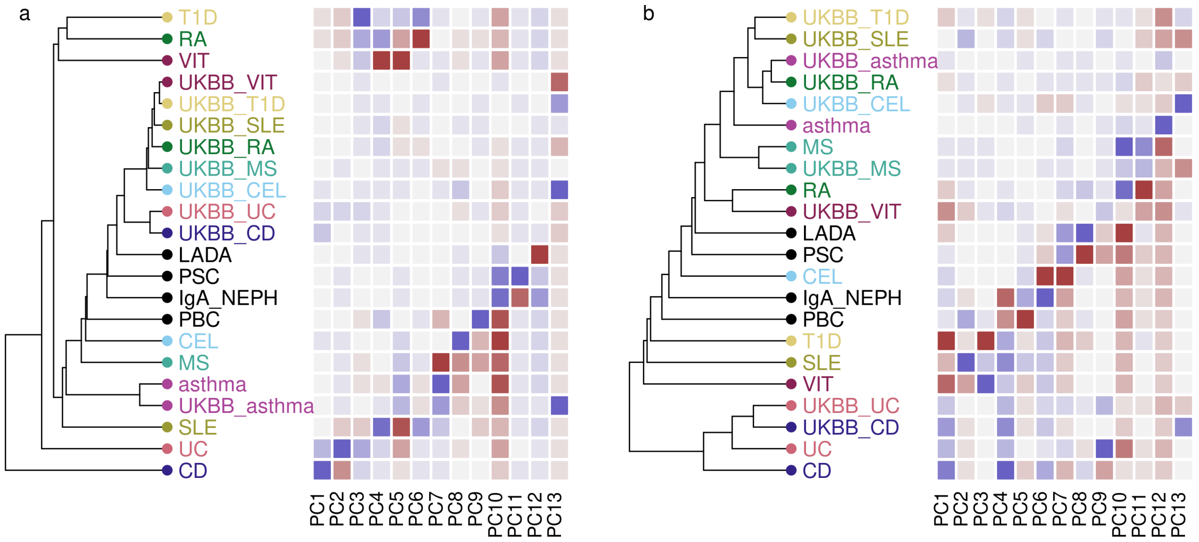


Supplementary Figure 1: Hierarchical clustering of basis diseases and their UKBB counterparts in **a** hard-thresholded, LD-thinned basis constructed using Z scores **b** hard-thresholded, LD-thinned basis constructed using $\hat{\beta}$ . Heatmaps indicate projected $\hat{\delta}$ for each disease on each component PC1-PC13, with grey indicating 0 (no difference from control), and darker shades of blue or magenta showing departure from controls in one direction or the other. GWAS datasets: T1D = type 1 diabetes, CEL= celiac disease, asthma, MS =multiple sclerosis, UC =ulcerative colitis, CD = Crohn's disease, RA =rheumatoid arthritis, VIT =vitiligo, SLE =systemic lupus erythematosus, PSC=primary sclerosing cholangitis, PBC=primary biliary cholangitis, LADA=latent autoimmune diabetes in adults, IgA_NEPH= IgA nephropathy. UKBB_ prefixed diseases correspond to self reported disease status in UK Biobank.

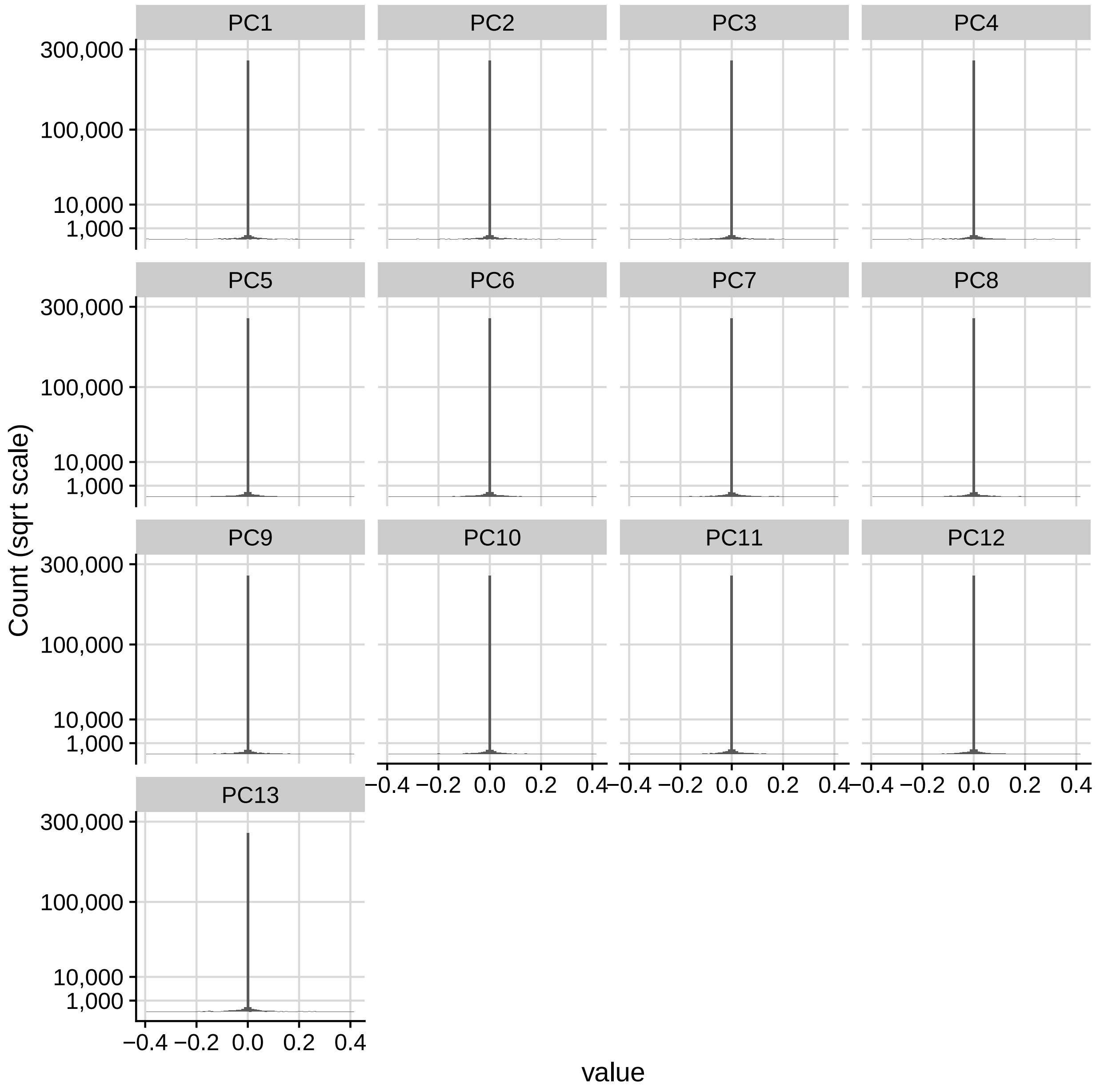


Supplementary Figure 2: Distributions of entries in the rotation matrix for each component PC1-PC13

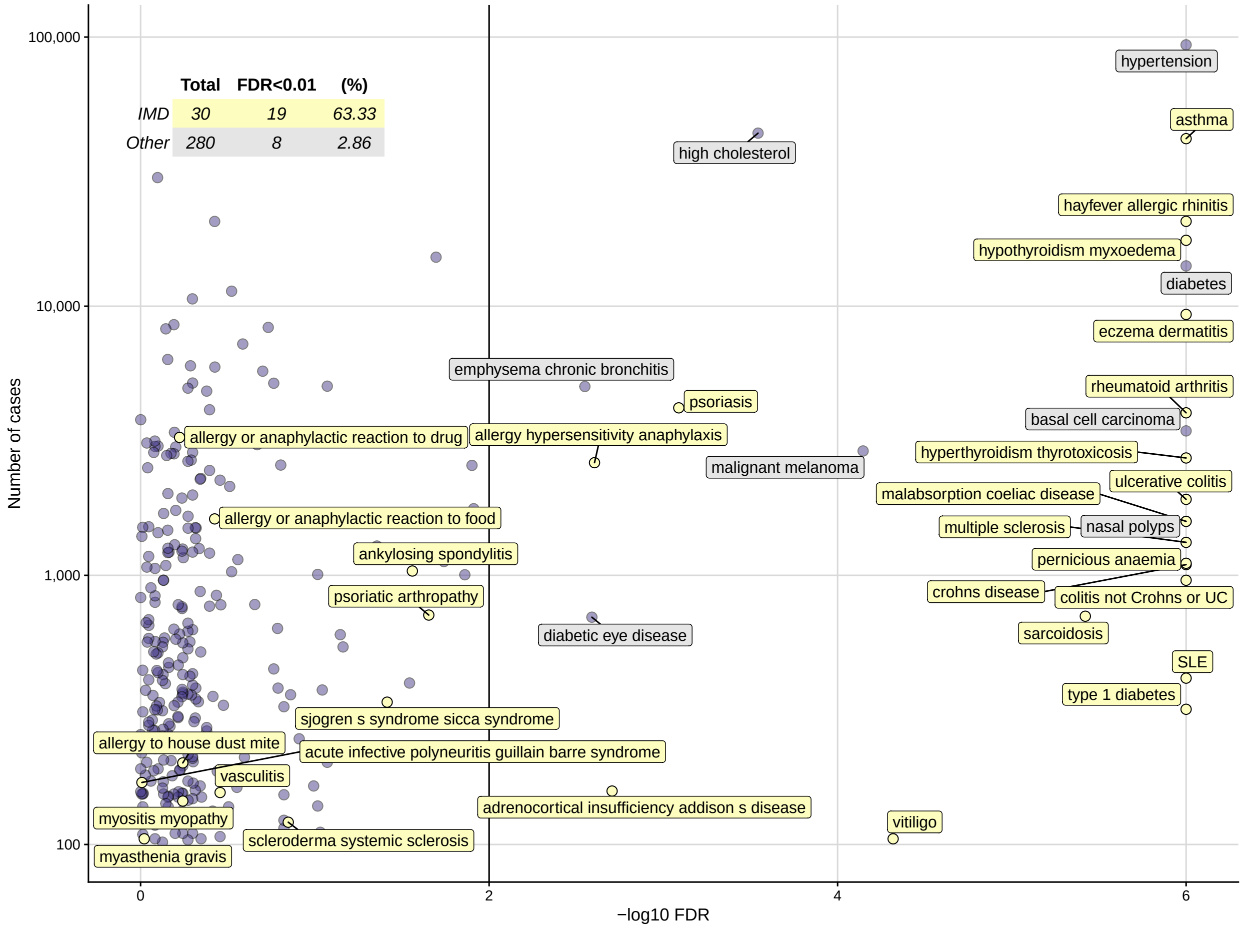


Supplementary Figure 3: IMD are enriched amongst significant (FDR < 1%) UKBB traits, with 63% of IMD even with few cases and < 3% of non-IMD traits significant. x axis is truncated at FDR=$10^{-6}$ for display.

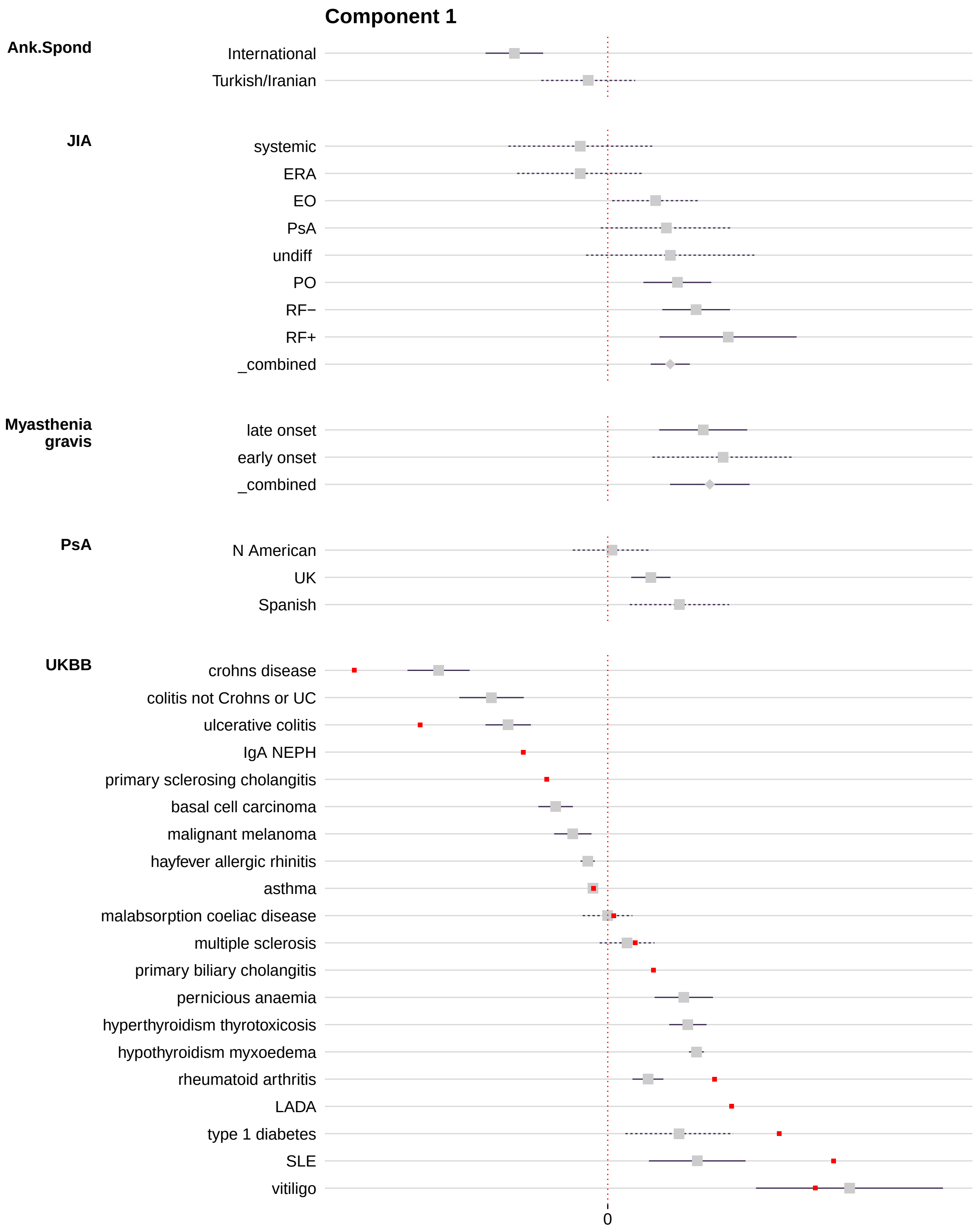


Supplementary Figure 4: Forest plot of component PC1 showing projected delta and 95% confidence interval (solid line = FDR < 1%, dashed line = FDR ≥ 1%). All IMD that are part of a trait group with at least one result significant at FDR < 1% are shown, together with any UKBB significant traits. IMD basis disease locations are shown in red.

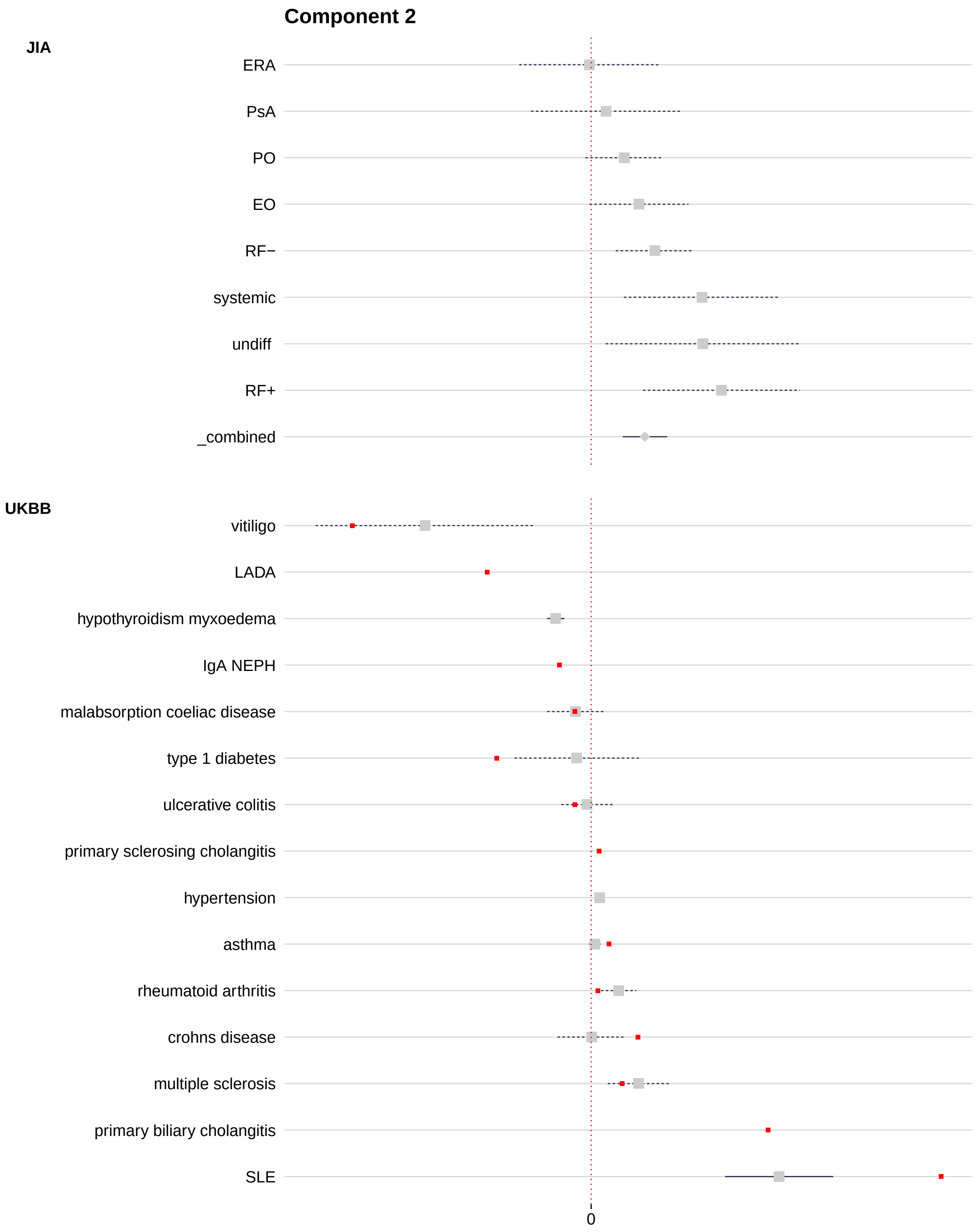


Supplementary Figure 5: Forest plot of component PC2 showing projected delta and 95% confidence interval (solid line = FDR $< 1\%$, dashed line = FDR $\geq 1\%$). All IMD that are part of a trait group with at least one result significant at FDR $< 1\%$ are shown, together with any UKBB significant traits. IMD basis disease locations are shown in red.

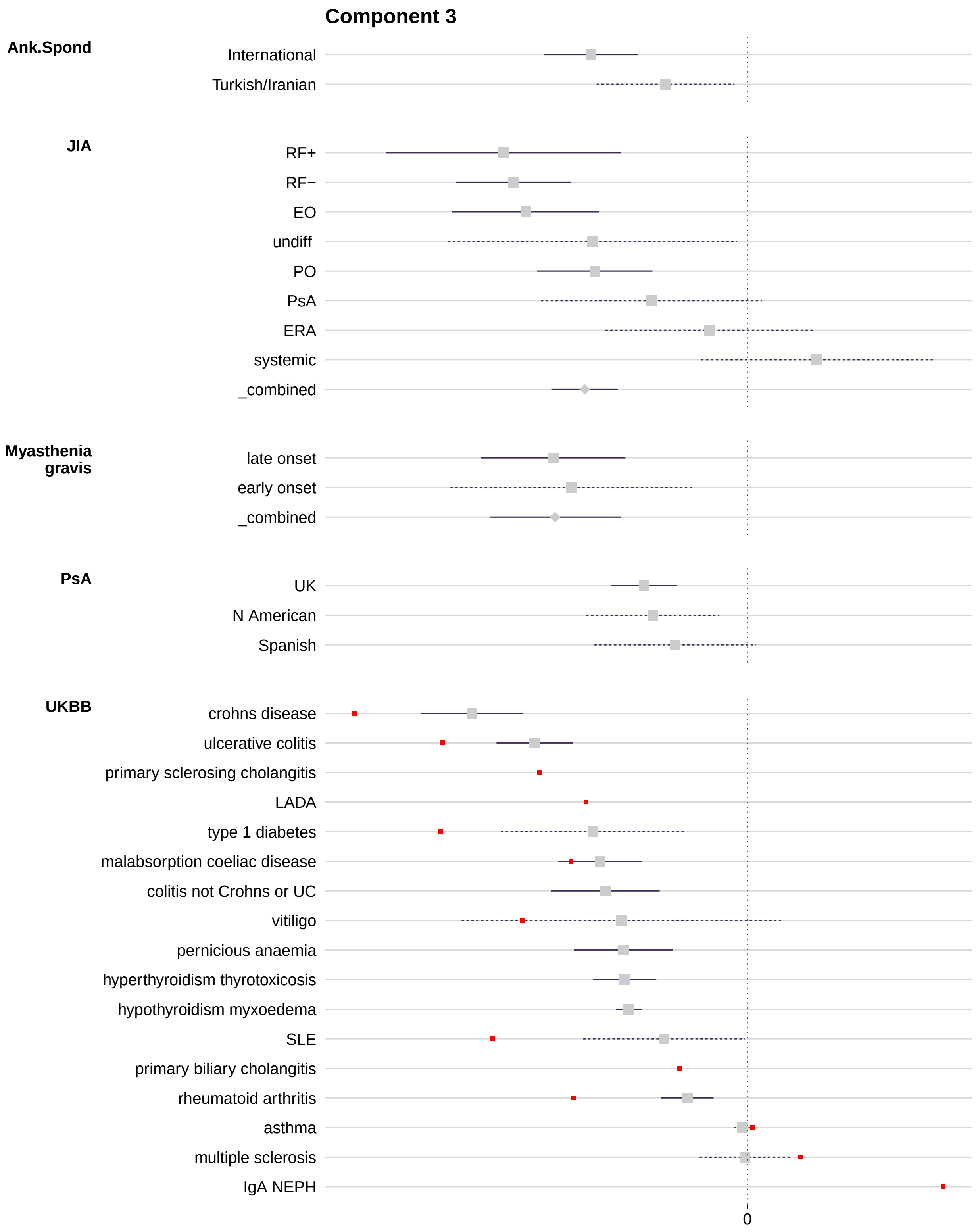


Supplementary Figure 6: Forest plot of component PC3 showing projected delta and 95% confidence interval (solid line = FDR $< 1\%$, dashed line = FDR $\geq 1\%$). All IMD that are part of a trait group with at least one result significant at FDR $< 1\%$ are shown, together with any UKBB significant traits. IMD basis disease locations are shown in red.

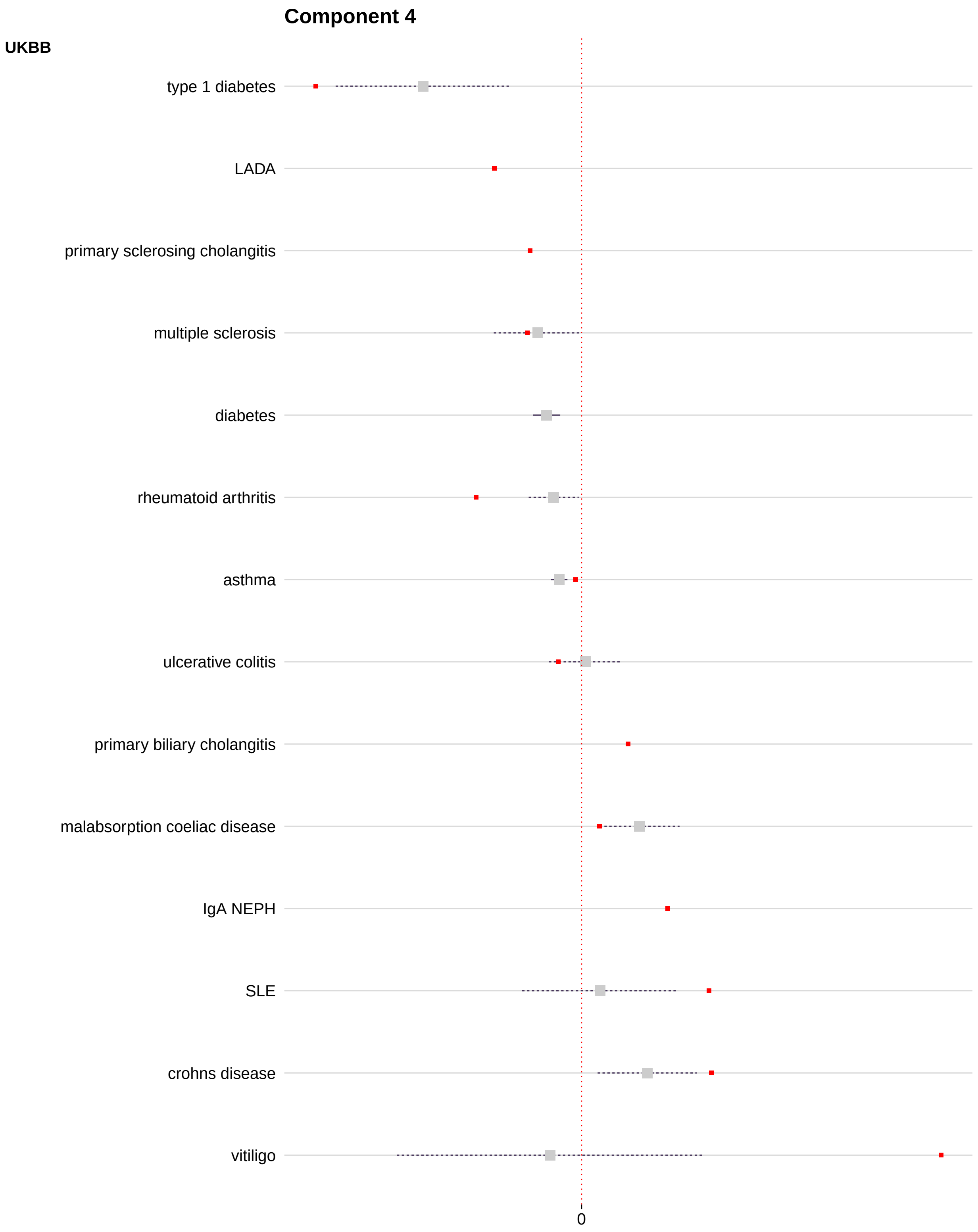


Supplementary Figure 7: Forest plot of component PC4 showing projected delta and 95% confidence interval (solid line = FDR $< 1\%$, dashed line = FDR $\geq 1\%$). All IMD that are part of a trait group with at least one result significant at FDR $< 1\%$ are shown, together with any UKBB significant traits. IMD basis disease locations are shown in red.

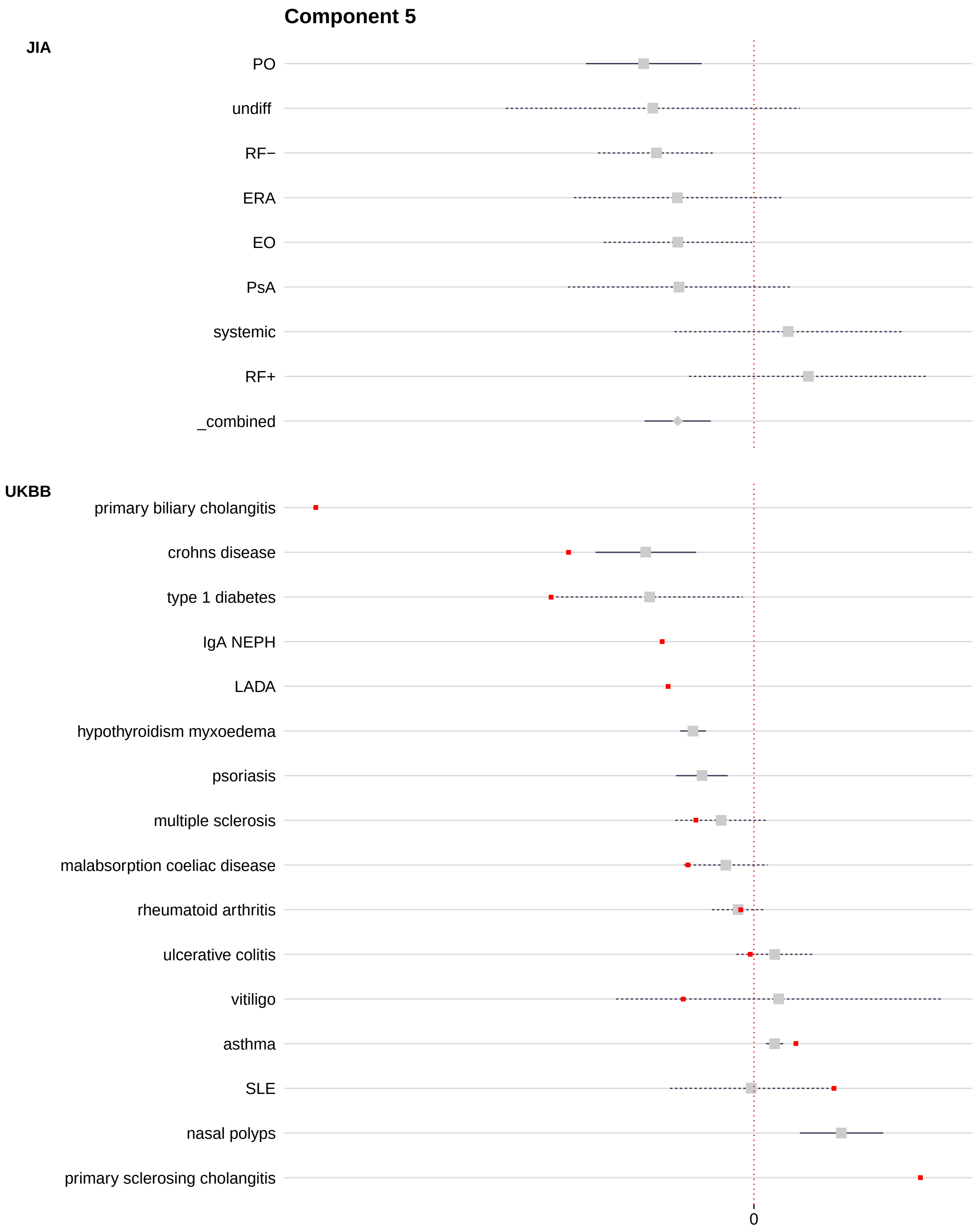


Supplementary Figure 8: Forest plot of component PC5 showing projected delta and 95% confidence interval (solid line = FDR $< 1\%$, dashed line = FDR $\geq 1\%$). All IMD that are part of a trait group with at least one result significant at FDR $< 1\%$ are shown, together with any UKBB significant traits. IMD basis disease locations are shown in red.

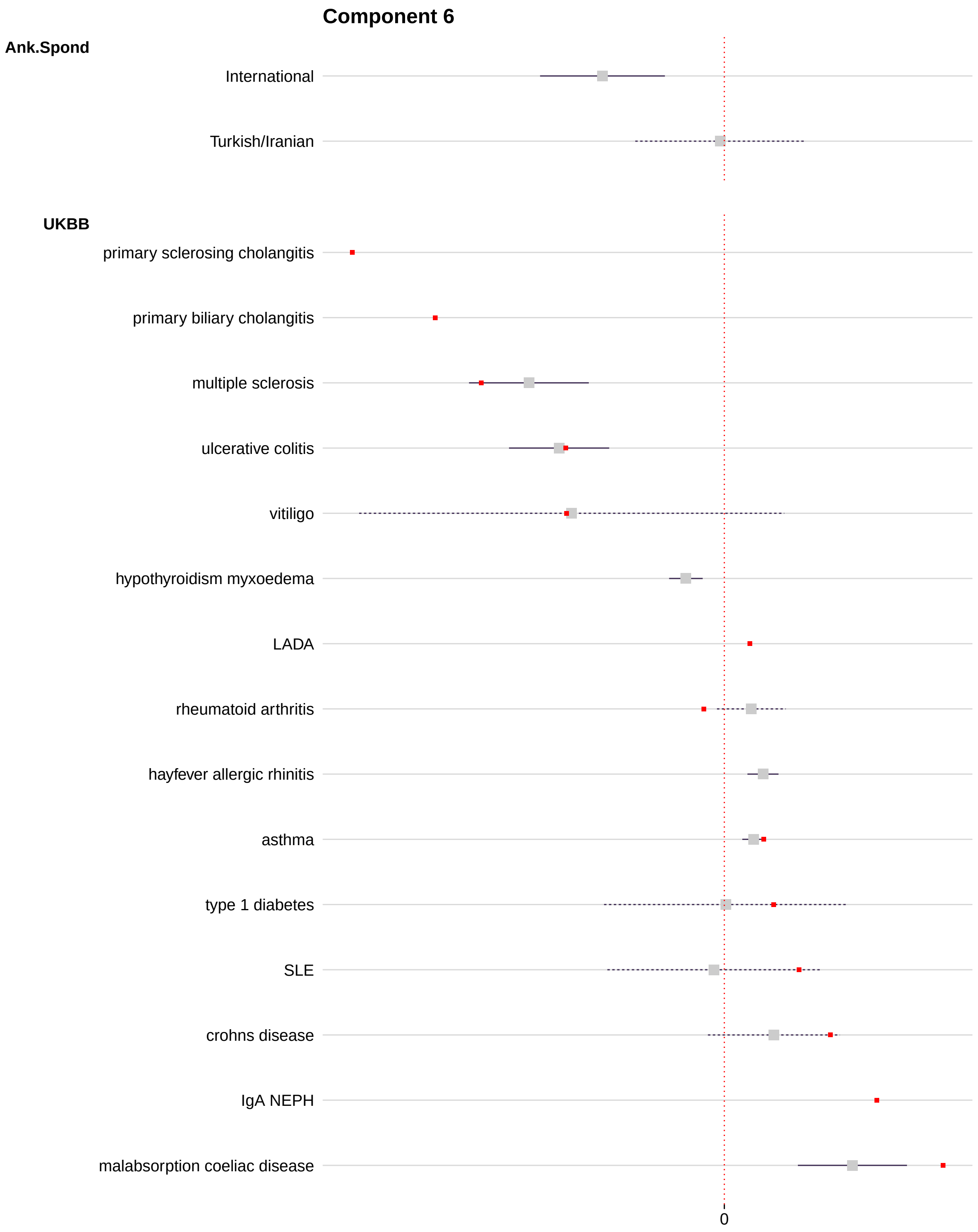


Supplementary Figure 9: Forest plot of component PC6 showing projected delta and 95% confidence interval (solid line = FDR $< 1\%$, dashed line = FDR $\geq 1\%$). All IMD that are part of a trait group with at least one result significant at FDR $< 1\%$ are shown, together with any UKBB significant traits. IMD basis disease locations are shown in red.

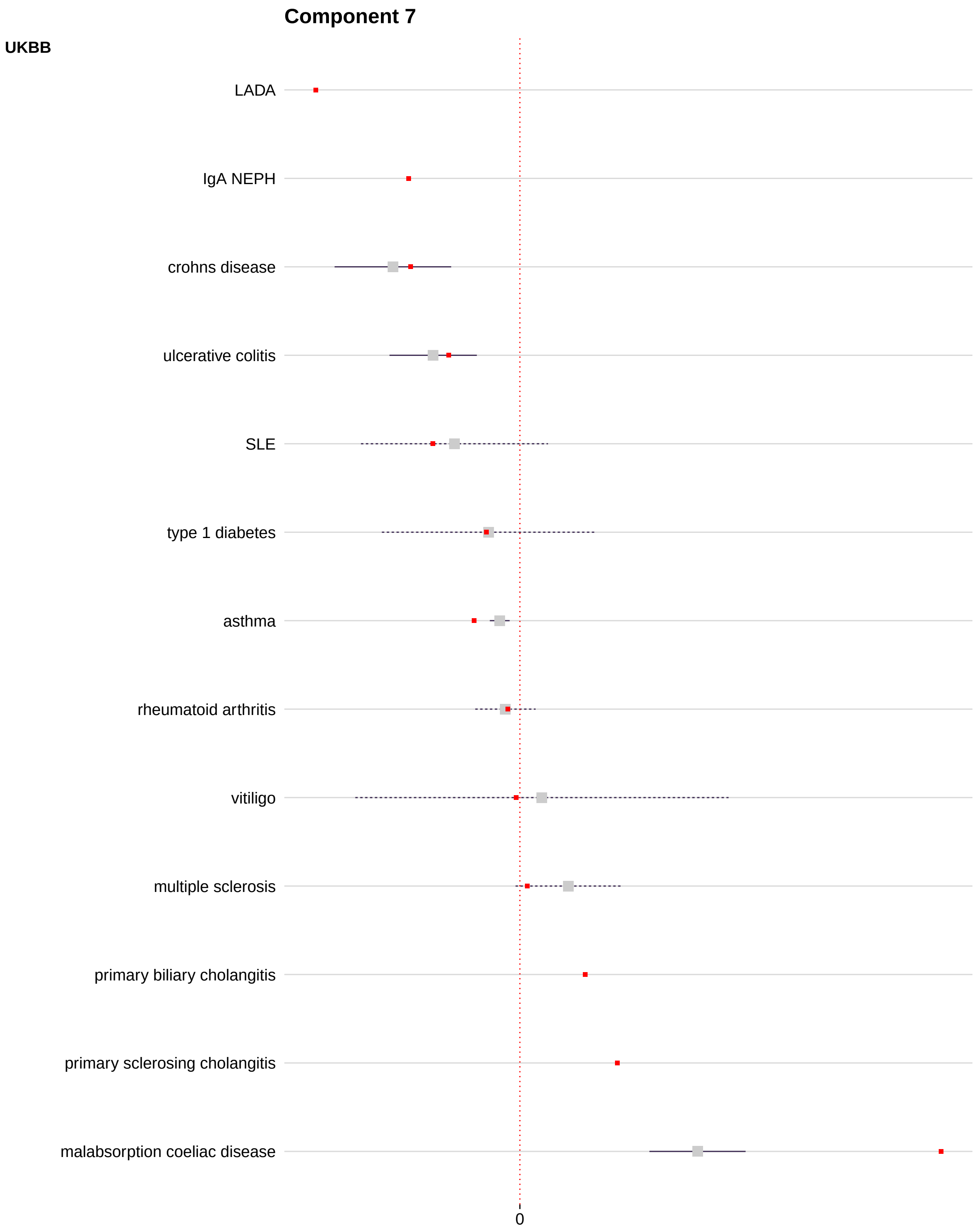


Supplementary Figure 10: Forest plot of component PC7 showing projected delta and 95% confidence interval (solid line = FDR $< 1\%$, dashed line = FDR $\geq 1\%$). All IMD that are part of a trait group with at least one result significant at FDR $< 1\%$ are shown, together with any UKBB significant traits. IMD basis disease locations are shown in red.

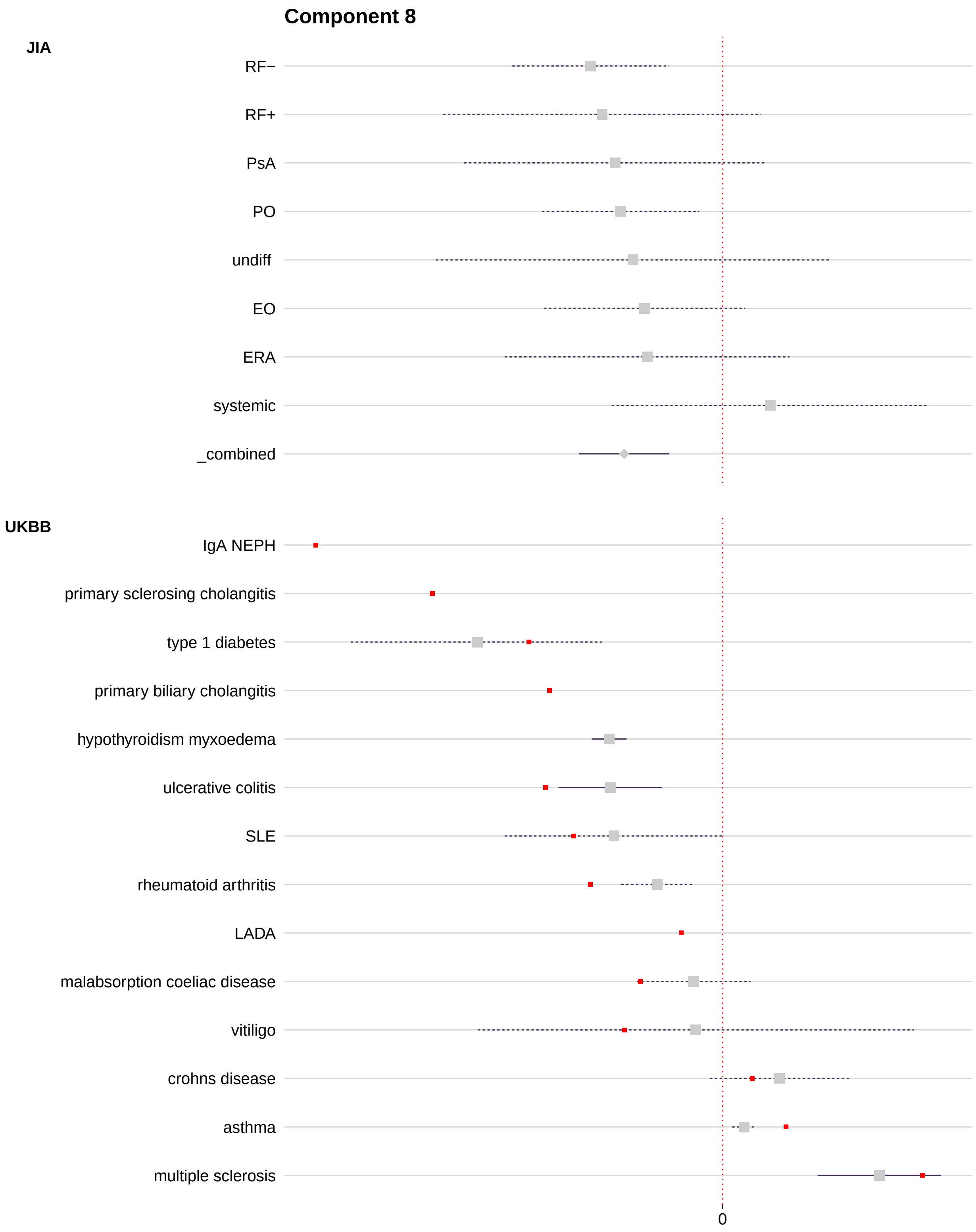


Supplementary Figure 11: Forest plot of component PC8 showing projected delta and 95% confidence interval (solid line = FDR < 1%, dashed line = FDR ≥ 1%). All IMD that are part of a trait group with at least one result significant at FDR < 1% are shown, together with any UKBB significant traits. IMD basis disease locations are shown in red.

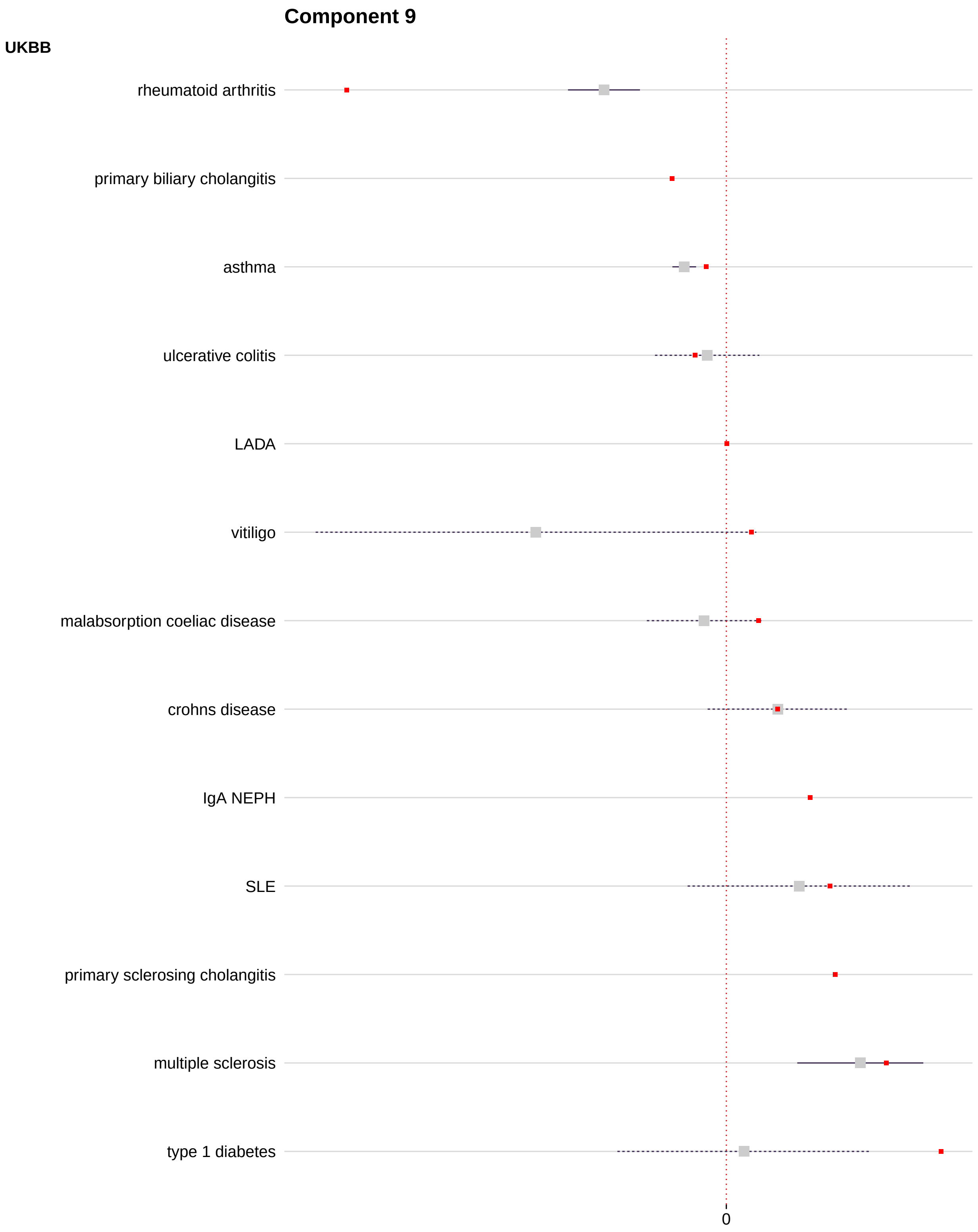


Supplementary Figure 12: Forest plot of component PC9 showing projected delta and 95% confidence interval (solid line = FDR $< 1\%$, dashed line = FDR $\geq 1\%$). All IMD that are part of a trait group with at least one result significant at FDR $< 1\%$ are shown, together with any UKBB significant traits. IMD basis disease locations are shown in red.

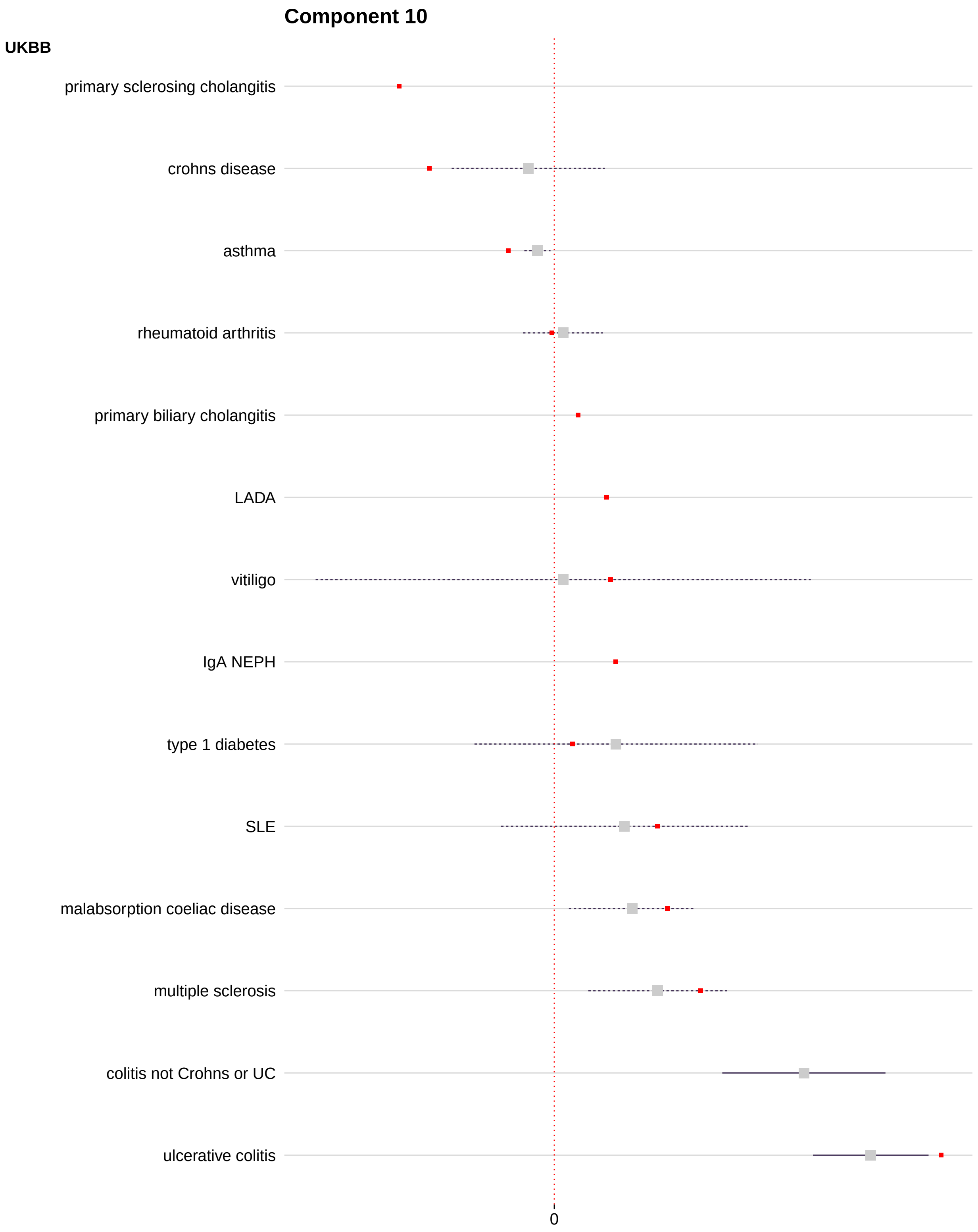


Supplementary Figure 13: Forest plot of component PC10 showing projected delta and 95% confidence interval (solid line = FDR < 1%, dashed line = FDR $\geq$ 1%). All IMD that are part of a trait group with at least one result significant at FDR < 1% are shown, together with any UKBB significant traits. IMD basis disease locations are shown in red.

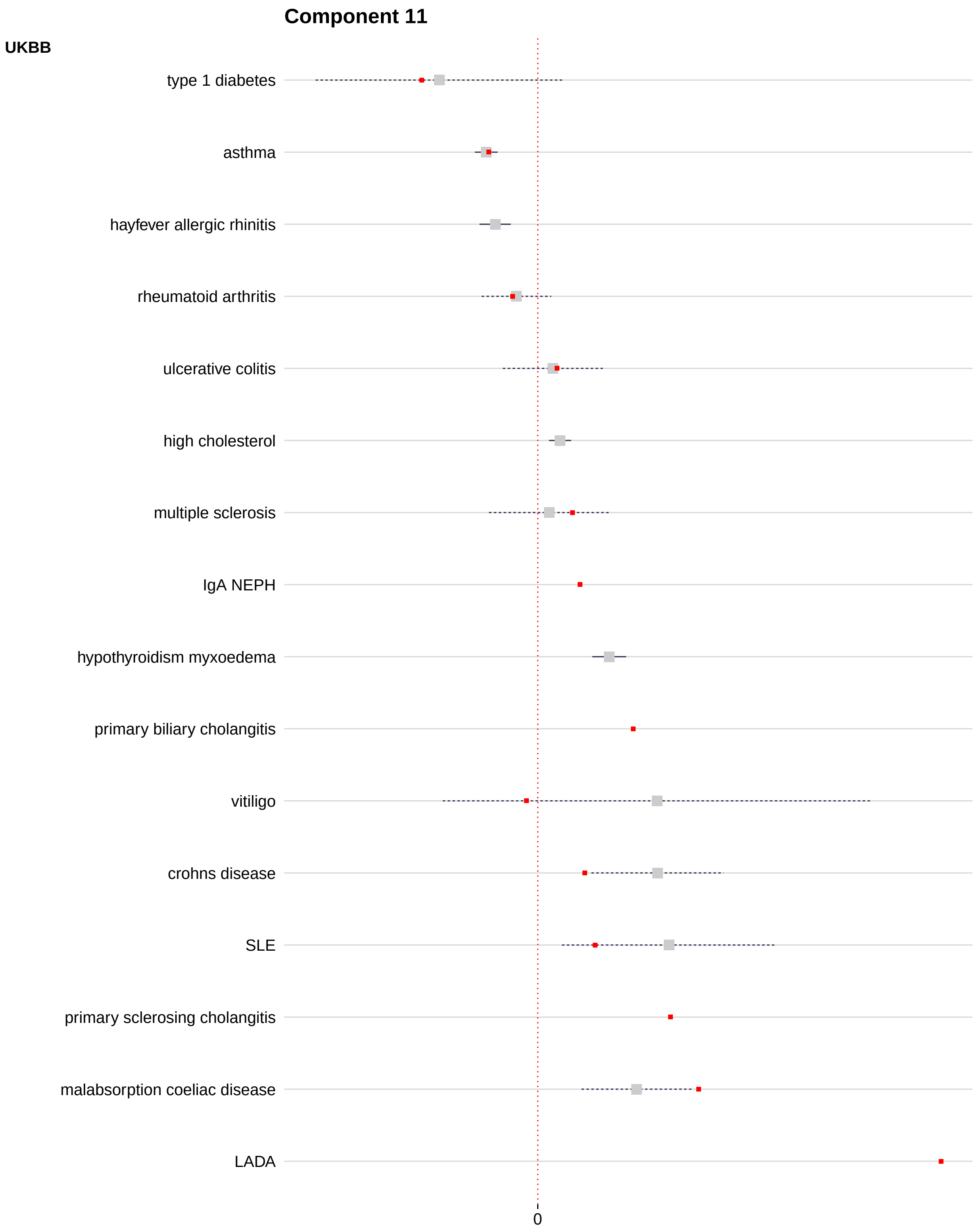


Supplementary Figure 14: Forest plot of component PC11 showing projected delta and 95% confidence interval (solid line = FDR $< 1\%$, dashed line = FDR $\geq 1\%$). All IMD that are part of a trait group with at least one result significant at FDR $< 1\%$ are shown, together with any UKBB significant traits. IMD basis disease locations are shown in red.

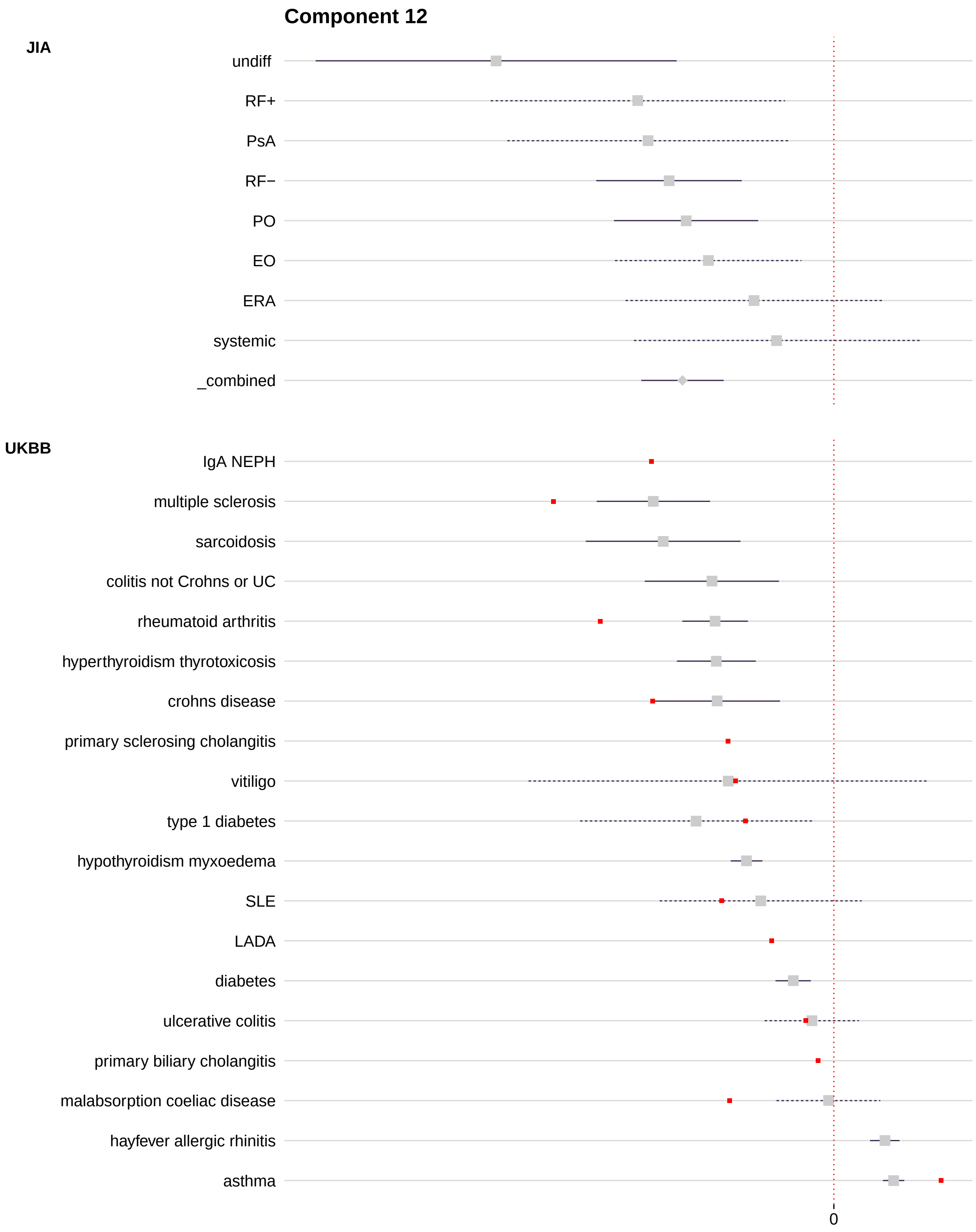


Supplementary Figure 15: Forest plot of component PC12 showing projected delta and 95% confidence interval (solid line = FDR $< 1\%$, dashed line = FDR $\geq 1\%$). All IMD that are part of a trait group with at least one result significant at FDR $< 1\%$ are shown, together with any UKBB significant traits. IMD basis disease locations are shown in red.

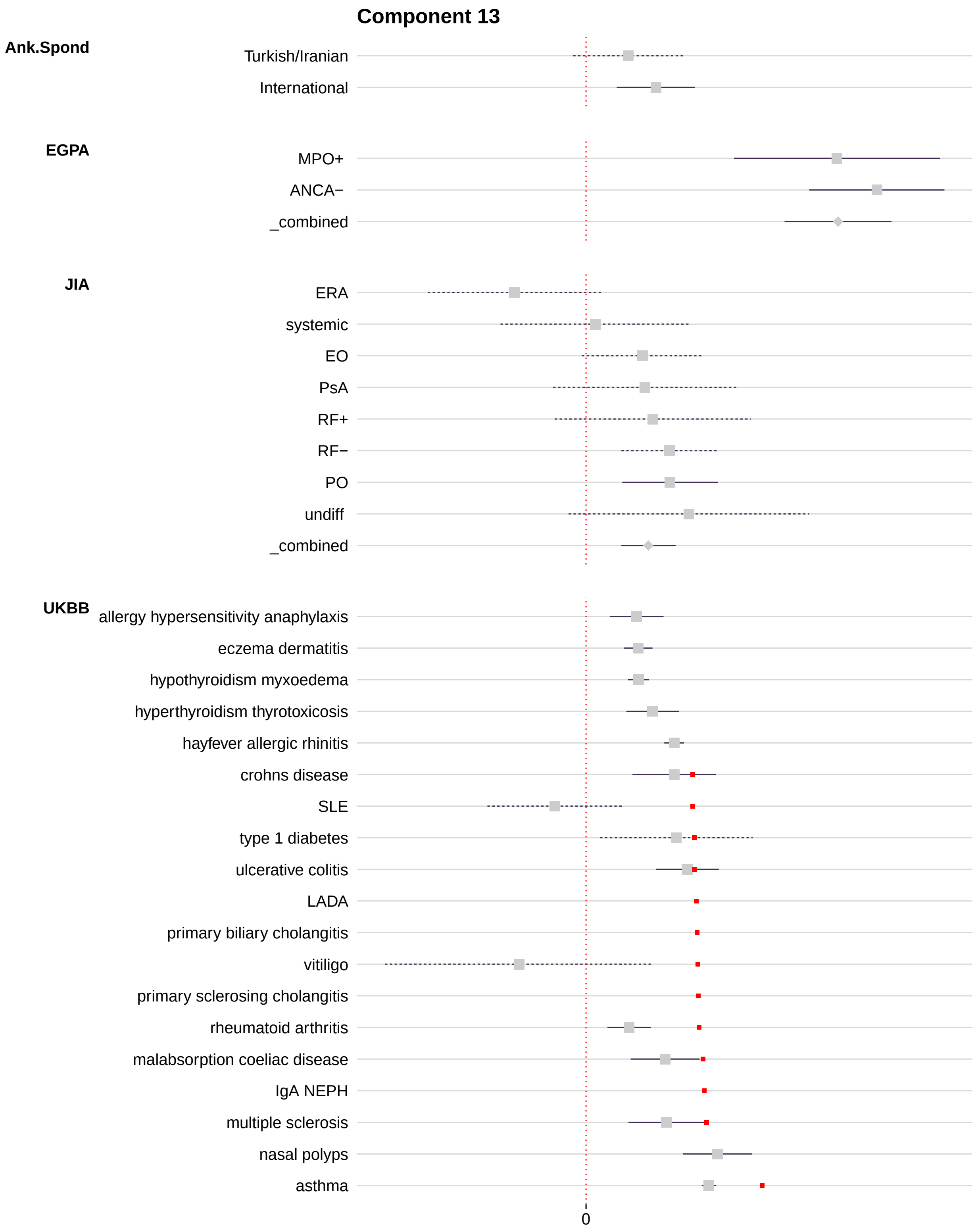


Supplementary Figure 16: Forest plot of component PC13 showing projected delta and 95% confidence interval (solid line = FDR $< 1\%$, dashed line = FDR $\geq 1\%$). All IMD that are part of a trait group with at least one result significant at FDR $< 1\%$ are shown, together with any UKBB significant traits. IMD basis disease locations are shown in red.

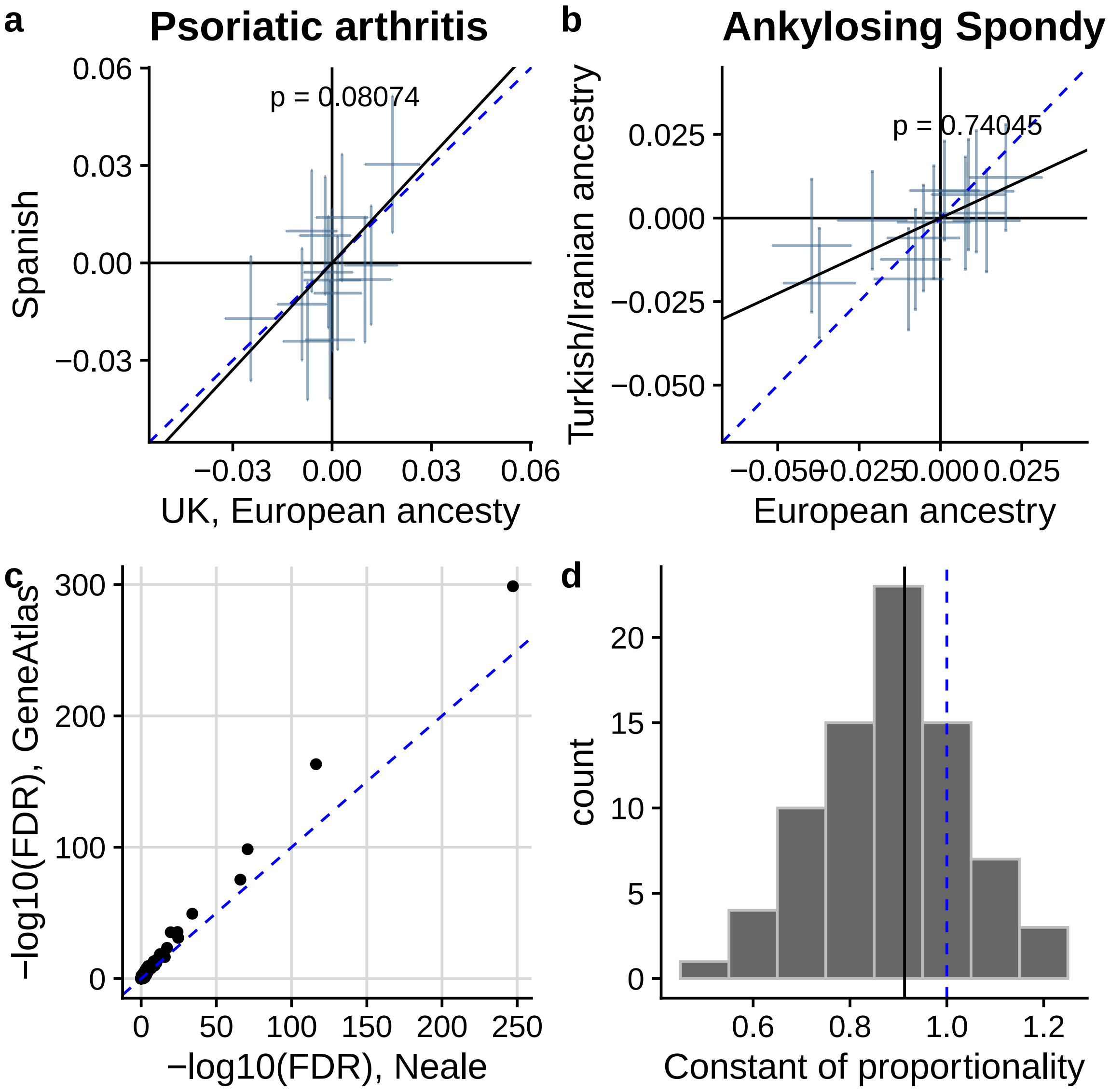


Supplementary Figure 17: Comparison of projects of the same traits from datasets with different ancestries. (a) in psoriatic arthritis, projections from a Spanish and UK (European ancestry) data set are shown with their 95% confidence intervals for each PC. Projections are very similar, with a constant of proportionality (black line) close to 1 (blue dashed line). The null hypothesis of proportionality is not rejected (p=0.08). (b) in ankylosing spondylitis, there is a more pronounced difference with an attenuation of the projections towards 0 in a Turkish/Iranian ancestry dataset compared to European ancestry, though the two remain proportional (p=0.74). In UKBB, Neale analysed only a white European subset while GeneAtlas includes additional non-European cases. Over 78 traits found in both datasets with similar case counts (within 10%), the additional sample size in GeneAtlas results in (c) generally more significant projections while (d) the constant of proportionality (GeneAtlas/Neale) is centred on 0.9.

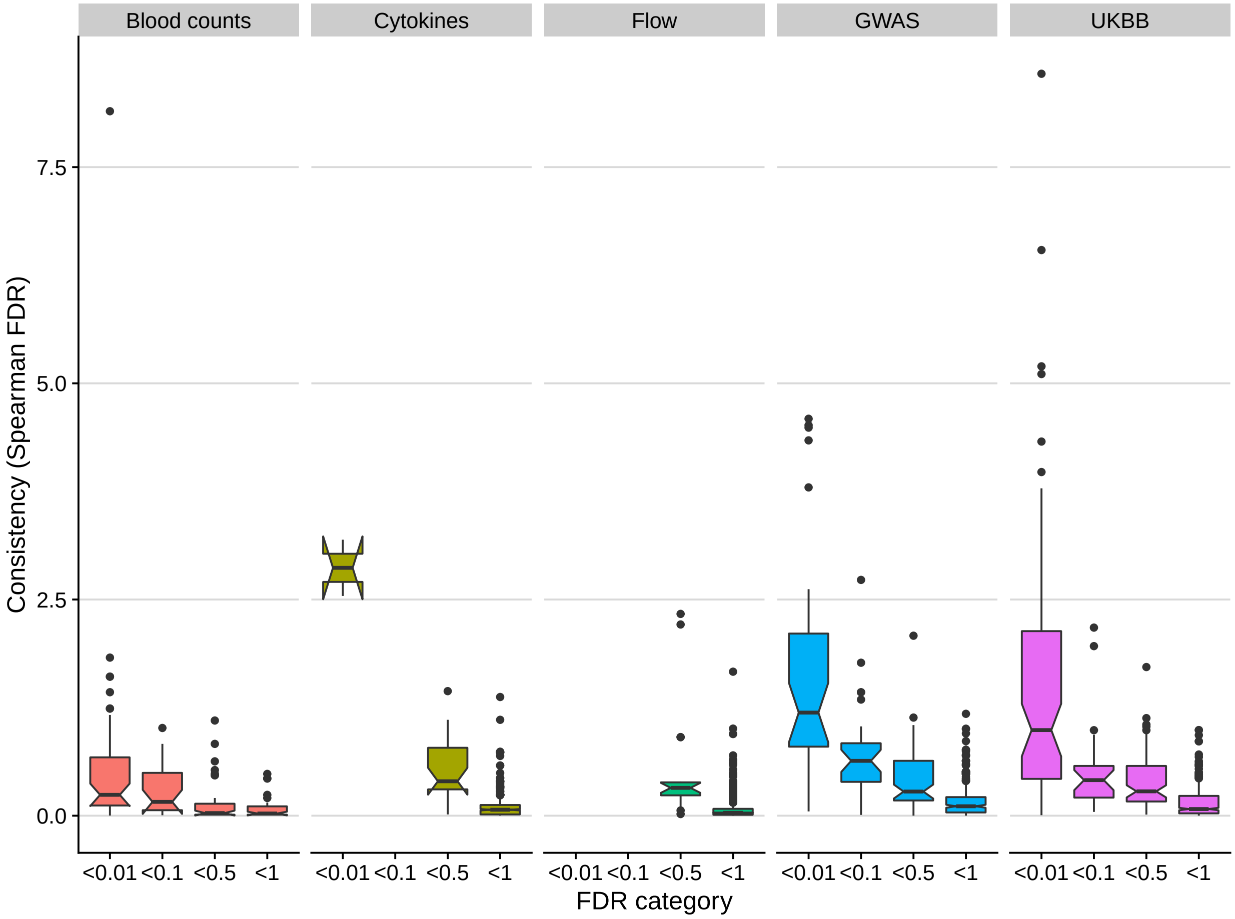


Supplementary Figure 18: For each group of traits, we compared the significance of Spearman correlation test of basis component SNP weights with trait $\beta$ (evidence for consistency, see Supplementary Note) with component FDR. We found traits with increasing component significance (smaller FDR) tended to also show more significant Spearman correlations, although the pattern was much weaker for blood cell counts despite more observations with small component FDRs. We subsequently filtered blood cell counts to count as significant only the outlying trait which was clearly significant by both component FDR and Spearman correlation, corresponding to eosinophil counts on PC13. Datasets are grouped by: blood cell counts,[21] cytokines,[23] flow cytometric immune cell counts,[22] GWAS datasets except UKBB, UKBB (Neale).

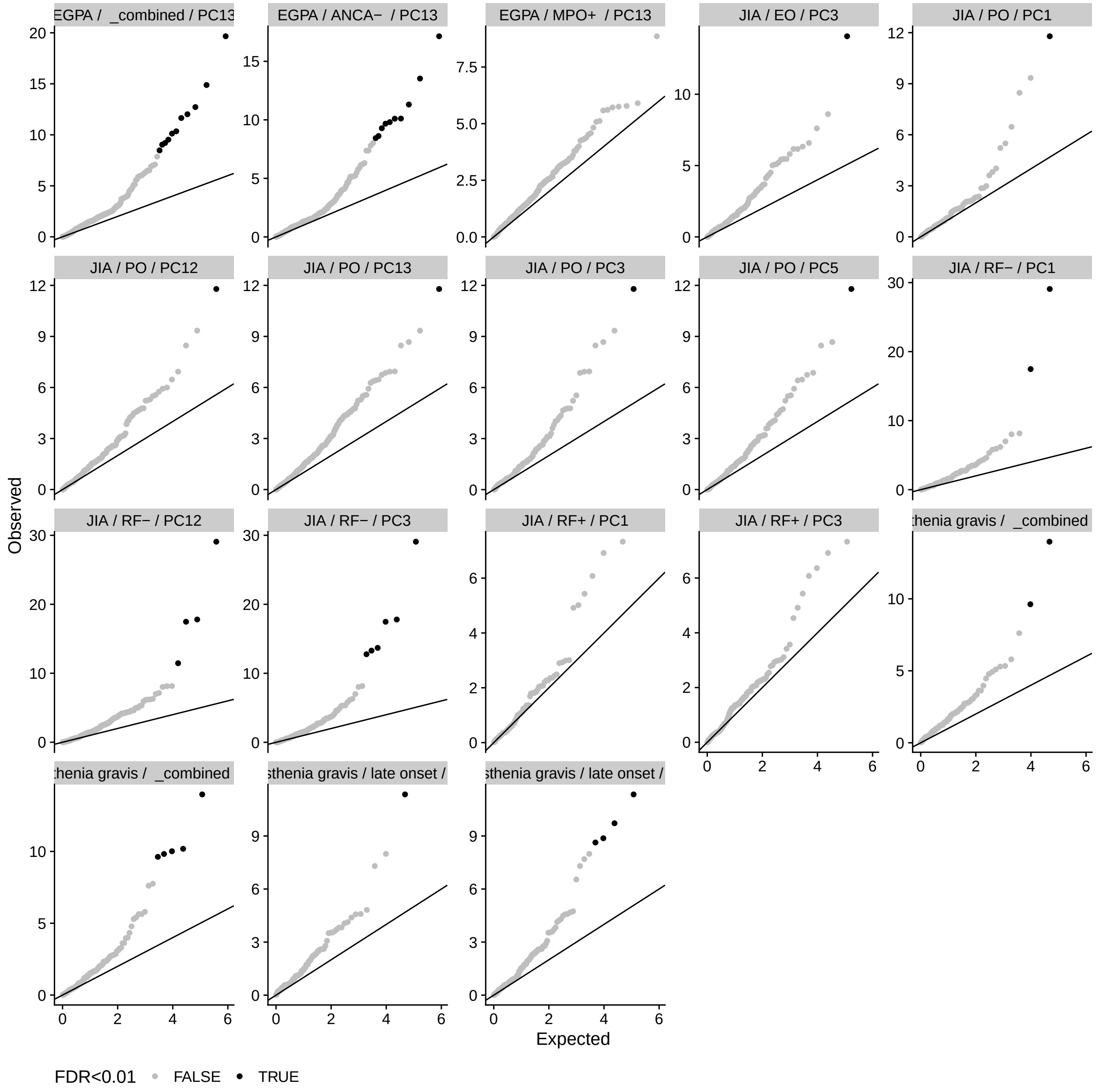


Supplementary Figure 19: QQ plots of p values for driver SNPs on trait-significant components showed a tendency for excess significant results. Points corresponding to FDR $< 1\%$ are highlighted in black, other points in gray. The solid line represents $y = x$.

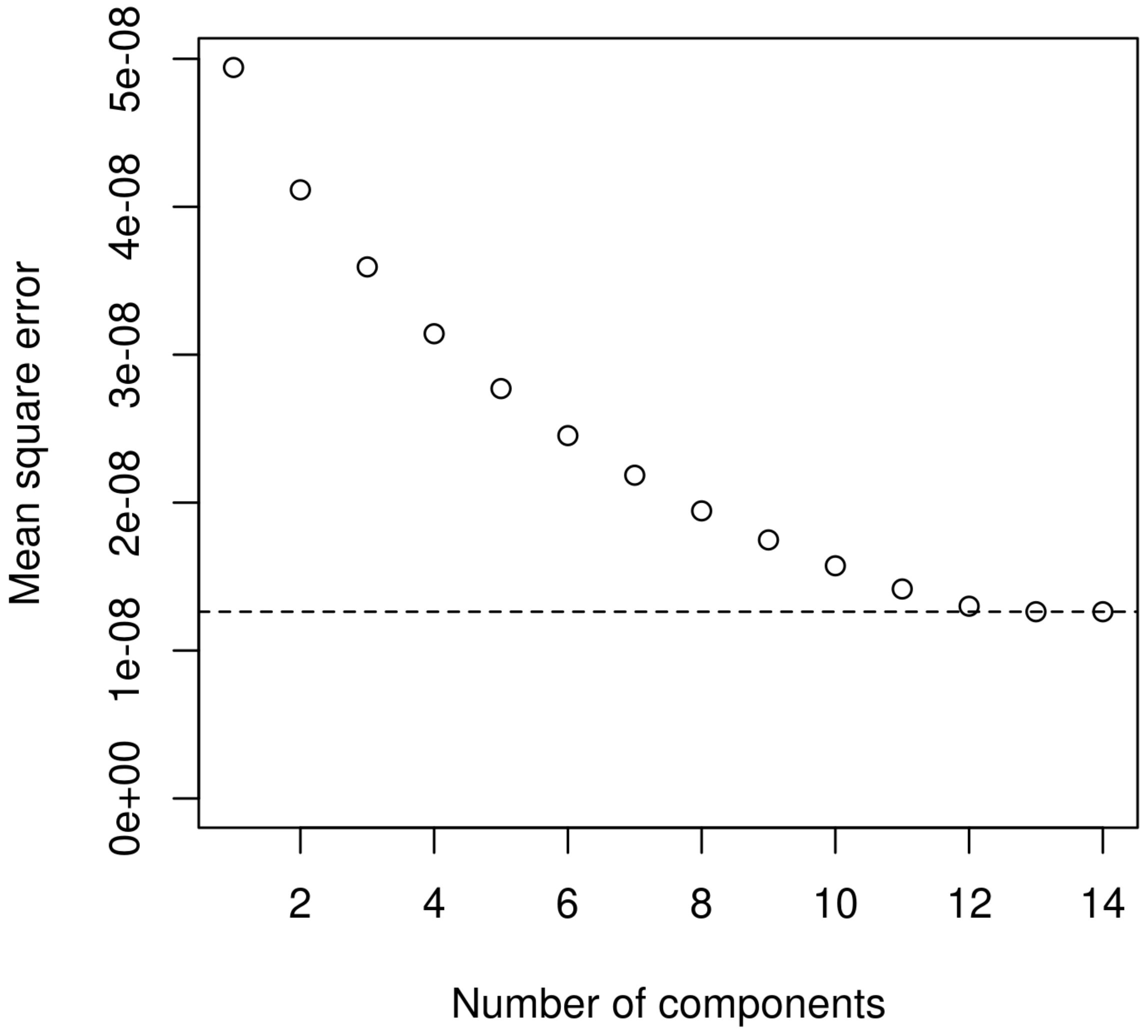


Supplementary Figure 20: Mean squared reconstruction error as the number of components used from the principal component decomposition increases from 1 to 14. The error is minimised with 13 or 14 components.
