## Supplementary Note for "Informed dimension reduction of clinically-related genome-wide association summary data characterises cross-trait axes of genetic risk"

### Characterisation of the genetic architecture of immune mediated disease through informed dimension reduction
### Supplementary Note

Oliver S Burren Guillermo Reales Limy Wong John Bowes James C Lee
Anne Barton Paul A Lyons Kenneth GC Smith Wendy Thomson
Paul DW Kirk Chris Wallace

October 2019

#### 1 Mathematical exposition of basis construction

Our input data are matrices of GWAS regression coefficients $\hat{\beta}_{ti}$ and their standard errors $\sigma^2_{ti}$ where $t = 1, \dots, T$ indexes traits, $i = 1, \dots, p$ indexes SNPs, and there are no missing values by design of picking only SNPs with complete data across all traits.

We first generate trait-specific weights for each SNP, using approaches developed for fine mapping. Wakefield[1] showed that the Bayes factor for SNP association could be approximated from GWAS summary statistics by

$$BF_{ti} = \frac{P(\hat{\beta}_{ti} | \hat{\beta}_{ti} \sim N(0, \sigma^2_{ti} + W)}{P(\hat{\beta}_{ti} | \hat{\beta}_{ti} \sim N(0, \sigma^2_{ti}))}$$

where $W$ is the variance of a prior on the true effect, and for which we set $W = 0.04$, a commonly adopted value (see, for example, [2]), corresponding to a prior belief that true odds ratios exceed 1.5 with a probability about 5%.

We assume the genome is partitioned into $R$ non-overlapping sets of SNPs, $\mathcal{R}_1, \dots, \mathcal{R}_R$, according to the location of recombination hotspots, such that each SNP belongs to exactly one region, and we write $i \in \mathcal{R}_r$ if the $i$-th SNP belongs to the $r$-th region. We use the notation $r(i)$ to denote the region to which SNP $i$ belongs; i.e. $r(i) = \mathcal{R}_j \iff i \in \mathcal{R}_j$. Then, under the assumption that at most one variant in a region $\mathcal{R}_r$ is causal, and that each such variant (if it exists) appears in the dataset, the posterior probability for each SNP $i \in \mathcal{R}_r$ to be causal is[3]

$$pp_{ti} = \frac{\pi BF_i}{(1 - m_r \pi) + \sum_{i \in \mathcal{R}_r} \pi BF_i}$$

where $m_r = |\mathcal{R}_r|$ is the number of SNPs in region $\mathcal{R}_r$ and we set the prior probability that any SNP in the dataset is causally associated with any trait to be $\pi = 10^{-4}$, corresponding to a prior belief that 1 in 10,000 SNPs across the genome is causal.[2] Although these trait-specific weights, $pp_{ti}$, were developed for fine-mapping, we emphasise that we are **not** fine mapping here, with only a subset of SNPs in our dataset. However, these weights are attractive because: (i) they tend to 0 for $\hat{\beta}_{ti}$ not distinguishable from

0; and (ii) they deal naturally with LD, since they are a function of the complete data likelihood for any recombination-hotspot defined region under the assumption the region contains a single causal variant.

We estimate the probability that the $r$-th region contains a causal association for the trait $t$ by summing across SNPs in that region

$$v_{tr} = \sum_{i \in \mathcal{R}_r} pp_{ti}$$

and use these to generate a final SNP weight which is a weighted average over $pp_{ti}$

$$w_i = \frac{\sum_t pp_{ti} v_{tr(i)}}{\sum_t v_{tr(i)}}$$

Each SNP also has a different variance in the population, $\sigma_i^2 = 2f_i(1-f_i)$ where $f_i$ is that SNP's minor allele frequency, and we use this to generate a $T \times p$ matrix of shrunk regression coefficients

$$(\hat{\gamma}_{ti}) = (w_i \beta_{ti} / \sigma_i)$$

to which we add a $T + 1^{th}$ row of zeroes, representing controls. A centred version of this matrix forms our final matrix for PCA decomposition

$$\boldsymbol{G}^C = (\hat{\gamma}_{ti}^C) = \left( \hat{\gamma}_{ti} - \frac{1}{T+1} \sum_t \hat{\gamma}_{ti} \right) = (\hat{\gamma}_{ti} - C_i)\,. \tag{1}$$

Standard PCA may be used to project $\boldsymbol{G}^C$ into a $T$-dimensional space by post-multiplying by the $p \times T$ projection matrix $Q$, whose columns are the first $T$ eigenvectors of $(\boldsymbol{G}^C)'\boldsymbol{G}^C$, to obtain

$$\boldsymbol{P} = \boldsymbol{G}^C \boldsymbol{Q}.$$

To project a new trait into the space, we require a vector of regression coefficients for that trait across the same set of SNPs, $\hat{\boldsymbol{\beta}}_0 = (\hat{\beta}_{0i}, i = 0, \dots, p)$. Typically these will be log OR for logistic regression, but they could also be from linear regression or any generalised linear model. Each $\hat{\beta}_{0i}$, has a corresponding estimate of variance $\nu_{0i}^2$, and the variance-covariance matrix of standardised $\hat{\boldsymbol{\beta}}_0$, $\boldsymbol{Z} = (\hat{\beta}_{0i}/\nu_{0i})$ is $\boldsymbol{\Sigma}$, the correlation matrix of genotypes indexed by SNPs in $\hat{\boldsymbol{\beta}}_0$ [4]. The trait is then projected onto the basis according to the linear function

$$\boldsymbol{P}_0 = (\boldsymbol{D}\hat{\boldsymbol{\beta}}_0 - \boldsymbol{C})'\boldsymbol{Q}$$

where $\boldsymbol{C} = (C_i, i = 1, \dots, p)$ is the vector of centres defined in (1), $\boldsymbol{D}$ is a diagonal matrix with entries $w_i/\sigma_i$ for each SNP $i = 0, \dots, p$. Then, defining $\boldsymbol{V}_0$ as the diagonal matrix with entries $\nu_{0i}$,

$$\begin{aligned} \text{var}(\hat{\boldsymbol{\beta}}_0) &= \boldsymbol{V}_0 \boldsymbol{\Sigma} \boldsymbol{V}_0 \\ \text{var}\, \boldsymbol{P}_0 &= \boldsymbol{Q}'\boldsymbol{D}\boldsymbol{V}_0\boldsymbol{\Sigma}\boldsymbol{V}_0\boldsymbol{D}\boldsymbol{Q} \end{aligned}$$

The null location in this space (i.e. the projection of a zero vector) is given by $X_0 = -\boldsymbol{C}'\boldsymbol{Q}$, so that the difference between the projection of the new trait and control is given by $\hat{\delta} = \boldsymbol{P}_0 - X_0$, which also has variance $\text{var}(\boldsymbol{P}_0)$ as only $\boldsymbol{P}_0$ is random. A statistic, $X^2_{overall}$, can then be defined as follows to allow us to test the null hypothesis that the estimand $\delta$ is zero across all components

$$X^2_{overall} = \hat{\delta}' \,\text{var}(\boldsymbol{P}_0)^{-1}\hat{\delta} \sim \chi^2_T \quad \text{if the null hypothesis is true.}$$

Similarly, we may define statistic $X^2_j$ to allow us to test the hypothesis that component $j$ of $\delta$ is zero:

$$X^2_j = \hat{\delta}[j]^2 / \,\text{var}(\boldsymbol{P}_0)[j,j] \sim \chi^2_1 \quad \text{if the null hypothesis is true.}$$

#### 2 Effect of SNP density on basis

Due to differential genotype coverage between input traits there is a tension between the number of traits and the density of variants that can be included in the basis. To investigate the dependence of the basis on variant density we created two shrinkage bases using six of the original input studies that used imputation: a ‘sparse’ basis using a subset of 265,887 SNPs used in the main text basis and a ‘dense’ basis using 4,623,330 SNPs (non-palindromic, MAF>0.01) common across the six studies. We found that despite the dense basis containing an order of magnitude more SNPs the PC scores for input traits were highly correlated between bases (Figure 1).

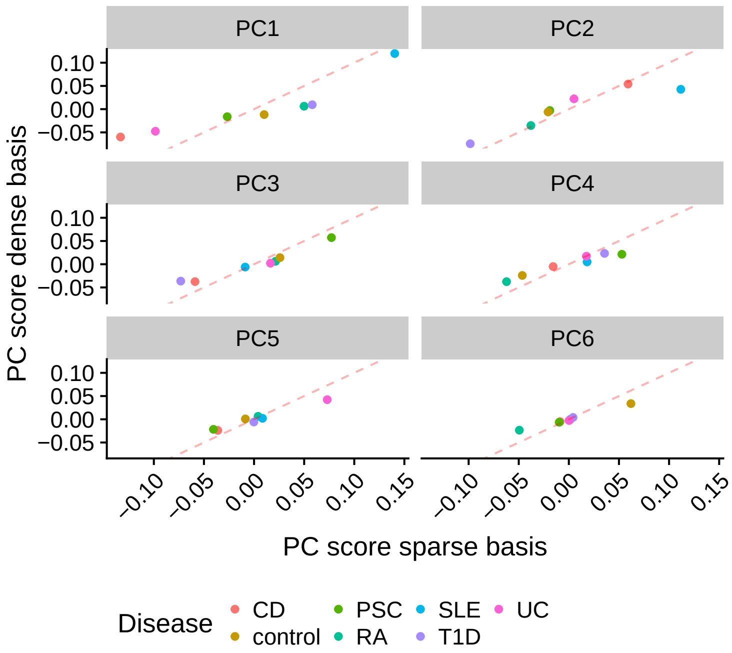


Figure 1: Comparison between ‘sparse’ and ‘dense’ bases for six input immune mediated diseases for which imputed summary statistics were available.

### 3 Missing variants in GWAS summary statistics for projection

Genotype coverage varied across the traits we projected onto the basis (Supplementary Table 3). By default, we set missing values to zero, and investigated the effect of this on resultant projections as follows. For each trait missing at least one basis SNP, we constructed an alternative basis using only SNPs present in all 13 basis traits and the trait to be projected. After projecting the target trait onto this alternative 'tailored' basis we assessed the difference in PC scores across components (Figure 2). We found in a majority of traits, PC scores were only nominally affected. Projections of NMO and PsA (Aterido) were substantially different in the tailored basis, mostly due to attenuated and thus conservative PC scores for the main basis. We concluded that these traits were sensitive to missing data and imputed summary statistics as described in Methods.

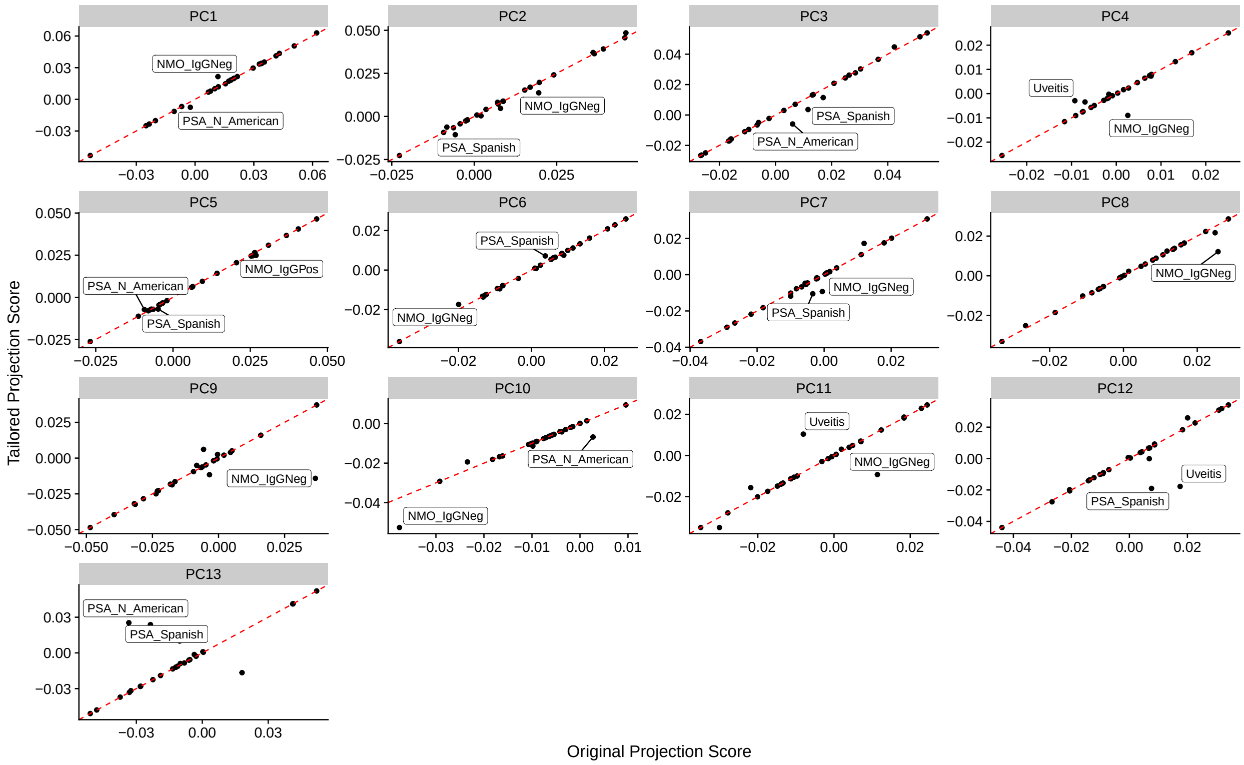


Figure 2: Comparison between original (missing $\hat{\beta}$ for the projected trait set to zero) and tailored basis (SNPs with missing $\hat{\beta}$ for the trait to be projected removed from all basis traits, a new basis created, and the trait projected) projection scores for traits in Supplementary Table 3. Traits with the largest deviations from $x = y$ are labelled and were subsequently imputed as described in Methods.

### 4 Validity of basis for different ancestry datasets

While our basis was created from predominantly European GWAS, there is an imperative to increase ancestry diversity in GWAS[5]. To assess whether the basis could be used for non-European ancestry, we compared projections of the same disease in different populations: studies of ankylosing spondylitis in Turkish and

Iranian[6], European[7] or British (UKBB) ancestry and PsA in individuals of Spanish or North American ancestry. We found projections were proportional across ancestries, though attenuated towards the null for the study with ancestry expected to be most distant to European, with ratios 0.52 (Turkish/Iranian AS compared to European AS) and 0.77 (Spanish PsA compared to North American PsA) (Supp Fig XX).

This was supported by a broader comparison of two releases of UKBB summary statistics. The Neale compendium used above was chosen because the traits analysed are not filtered on case numbers, allowing us to consider IMD with small numbers of cases. Whilst the Neale compendium focuses on the European subset of UKBB (n approx = 360,000) an alternative resource, GeneAtlas, (http://geneatlas.roslin.ed.ac.uk/)(Canela-Xandri, Rawlik, and Tenesa 2018) uses all available UKBB subjects (n approx= 452,000), and a linear mixed model to adjust for population stratification. GeneAtlas projections were generally proportional to Neale (no significant deviation from proportionality identified) but attenuated (median ratio=0.89). This suggests the mix of non-European samples leads to an attenuation of signal compared to European-only. However, the larger sample size in GeneAtlas compensated for the attenuation such that GeneAtlas results showed a tendency to greater significance (Supplementary Figure XX - suppfig-ancestry.pdf). Overall, this suggests that results projecting non-European or GWAS studies on to a European basis may reduce power for the same sample size, but does not lead to invalid results.

### References


[1] Jon Wakefield. Bayes factors for genome-wide association studies: comparison with P -values. *Genet. Epidemiol.*, 33(1):79–86, January 2009.

[2] Claudia Giambartolomei, Damjan Vukcevic, Eric E Schadt, Lude Franke, Aroon D Hingorani, Chris Wallace, and Vincent Plagnol. Bayesian test for colocalisation between pairs of genetic association studies using summary statistics. *PLoS Genet.*, 10(5):e1004383, May 2014.

[3] The Wellcome Trust Case Control Consortium, Julian B Maller, Gilean McVean, Jake Byrnes, Damjan Vukcevic, Kimmo Palin, Zhan Su, Joanna M M Howson, Adam Auton, Simon Myers, Andrew Morris, Matti Pirinen, Matthew A Brown, Paul R Burton, Mark J Caulfield, Alastair Compston, Martin Farrall, Alistair S Hall, Andrew T Hattersley, Adrian V S Hill, Christopher G Mathew, Marcus Pembrey, Jack Satsangi, Michael R Stratton, Jane Worthington, Nick Craddock, Matthew Hurles, Willem Ouwehand, Miles Parkes, Nazneen Rahman, Audrey Duncanson, John A Todd, Dominic P Kwiatkowski, Nilesh J Samani, Stephen C L Gough, Mark I McCarthy, Panagiotis Deloukas, and Peter Donnelly. Bayesian refinement of association signals for 14 loci in 3 common diseases. *Nat. Genet.*, 44(12):1294–1301, October 2012.

[4] Oliver S Burren, Hui Guo, and Chris Wallace. VSEAMS: a pipeline for variant set enrichment analysis using summary GWAS data identifies IKZF3, BATF and ESRRA as key transcription factors in type 1 diabetes. *Bioinformatics*, 30(23):3342–3348, December 2014.

[5] Giorgio Sirugo, Scott M Williams, and Sarah A Tishkoff. The Missing Diversity in Human Genetic Studies. *Cell*, 177(1):26–31, March 2019.

[6] Zhixiu Li, Servet Akar, Handan Yarkan, Sau Kuen Lee, Pınar Çetin, Gerçek Can, Gökce Kenar, Fernur Çapa, Omer Nuri Pamuk, Yavuz Pehlivan, Katie Cremin, Erika De Guzman, Jessica Harris, Lawrie Wheeler, Ahmadreza Jamshidi, Mahdi Vojdanian, Elham Farhadi, Nooshin Ahmadzadeh, Zeynep Yüce, Ediz Dalkılıç, Dilek Solmaz, Berrin Akın, Salim Dönmez, İsmail Sarı, Paul J Leo, Tony J Kenna, Fatos Önen, Mahdi Mahmoudi, Matthew A Brown, and Nurullah Akkoc. Genome-wide association study in Turkish and Iranian populations identify rare familial Mediterranean fever gene (MEFV) polymorphisms associated with ankylosing spondylitis. *PLoS Genet.*, 15(4):e1008038, April 2019.

[7] Australo-Anglo-American Spondyloarthritis Consortium (TASC), John D Reveille, Anne-Marie Sims, Patrick Danoy, David M Evans, Paul Leo, Jennifer J Pointon, Rui Jin, Xiaodong Zhou, Linda A Bradbury, Louise H Appleton, John C Davis, Laura Diekman, Tracey Doan, Alison Dowling, Ran Duan, Emma L Duncan, Claire Farrar, Johanna Hadler, David Harvey, Tugce Karaderi, Rebecca Mogg, Emma Pomeroy, Karena Pryce, Jacqueline Taylor, Laurie Savage, Panos Deloukas, Vasudev Kumanduri, Leena Peltonen, Sue M Ring, Pamela Whittaker, Evgeny Glazov, Gethin P Thomas, Walter P Maksymowych, Robert D Inman, Michael M Ward, Millicent A Stone, Michael H Weisman, B Paul Wordsworth, and Matthew A Brown. Genome-wide association study of ankylosing spondylitis identifies non-MHC susceptibility loci. *Nat. Genet.*, 42(2):123–127, February 2010.
